# Light Martini water accelerates sampling in coarse-grained molecular dynamics simulations

**DOI:** 10.64898/2026.08.03.741232

**Authors:** Aladdin Elgendy, André P. Zeipelt, Lars V. Schäfer

## Abstract

Molecular dynamics (MD) simulations of slow biomolecular processes, such as exploration of the conformational ensembles of intrinsically disordered proteins (IDPs), are computationally demanding. Although coarse-grained (CG) models can substantially speed up the simulations compared to all-atom MD, the sampling challenge can still be significant for large systems and long time scales. Here, we present *light Martini water*, a low-viscosity water model that accelerates sampling in MD simulations with the Martini CG force field. We systematically reduced the mass of the Martini water beads and verified stable, accurate integration of the equations of motion with 20 fs time steps, as typically used in Martini simulations. *Light Martini water* has a reduced mass of 20 amu (compared to 72 amu in the standard water model), yielding up to a 2.68-fold increase in the sampling rate of IDP chain reconfiguration in water and a 16 % increase in the lateral diffusion of lipids in a POPC bilayer. Equilibrium properties remained unaffected by the mass scaling, and the speedup was achieved without compromising simulation accuracy. The water model is trivial to implement, has no computational overhead, and should be universally applicable to Martini simulations.

## I. INTRODUCTION

Biomolecular systems such as proteins, lipid membranes, and nucleic acids operate through complex, dynamic processes that span a wide range of spatial and temporal scales. Understanding the conformational landscapes and functional mechanisms of these systems at atomic resolution is a central goal of structural biology and biophysics. Molecular dynamics (MD) simulations provide a powerful computational framework to achieve this, offering the ability to model the time evolution of biomolecular systems in great detail. Despite their success and wide adoption, MD simulations face a fundamental limitation: the sampling problem. While biologically relevant processes may occur on microsecond to millisecond time scales (or even slower), conventional MD simulations are typically restricted to time scales of up to a few microseconds, even on modern high-performance computing platforms. The computational expense of MD simulations scales linearly with the number of integration steps, which are defined by the integration time step Δ*t*. This time step has to be substantially smaller than the time scale of the fastest motions in the molecular system to ensure numerical stability and avoid catastrophic crashes between particles.

To accelerate sampling, two pragmatic levers are widely used in all-atom MD, both aiming to achieve larger particle displacements per integration time step. One is hydrogen mass repartitioning (HMR), which shifts mass from heavy atoms to bonded hydrogens and enables a time step increase from 2 fs to 4 fs – with constraints – while leaving equilibrium configuration averages essentially unchanged.^1–3^ Another approach was recently presented by Rosas Jiménez et al., who used mass repartitioning in conjunction with mass scaling to develop a water model with faster self-diffusion that enables a roughly twofold acceleration in sampling efficiency while keeping the 2 fs time step.^4^

The idea of increasing sampling efficiency in molecular simulations by modifying the particle masses dates back to the 1970s, with Bennett being among the first to demonstrate its use in an actual computer simulation (of a Lennard-Jones polymer, in that case).^5^ Walser et al. showed that water viscosity can be decreased by reducing the water masses^6^ and that the solvent viscosity itself does not influence equilibrium properties of proteins, i. e., the average structure and its fluctuations.^7^ Provided that the associated motions involve friction with the solvent, a lower solvent viscosity should result in a speedup of (bio-)chemical processes in the condensed phase.^8–11^ Importantly, mass scaling should not affect equilibrium configurational properties, because the partition function factorizes into a product of a momentum integral and a position integral, of which the former cancels out in the expression for the canonical ensemble average:

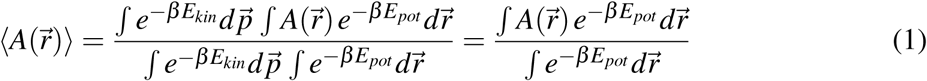

Here, *A*(*r⃗*) is the observable of interest, which only depends on the particle positions *r⃗* and not on their momenta 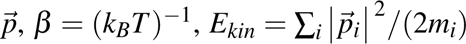 is the total kinetic energy, and *E_pot_* the total potential energy, which is independent of the particle masses.

Furthermore, the numerical integration of Newton’s equations of motion is invariant under mass scaling, provided that mass and time step are scaled by factors *s* and √*s*, respectively (*m*′ = *s* · *m* and Δ*t*′ = √*s* · Δ*t*). This also demonstrates that a uniform scaling of all masses in the system is equal to a rescaling of system time^1^ and would therefore not improve the sampling efficiency. The TIP3P-F water model developed by Rosas Jiménez et al.^4^ combines mass scaling (to accelerate diffusion) with mass repartitioning (to compensate for time-integration instabilities introduced by the mass scaling). Their model is based on the widely used three-site water model TIP3P. Rosas Jiménez et al. repartitioned 2 amu from the oxygen to each hydrogen atom and scaled the resulting masses by a factor of approximately 16, yielding a hydrogen mass of 0.186 amu and an oxygen mass of 0.744 amu. Compared to normal TIP3P water, the self-diffusion of TIP3P-F water is faster by a factor of approximately 3.3, which enhances the conformational sampling of the investigated biomolecules without substantially affecting structural and thermodynamic properties. They observed a roughly twofold sampling speedup for the end-to-end distance and backbone dihedral angle order parameters of small peptides and nucleotides and for the radius of gyration of the protein ubiquitin. Furthermore, a lipid diffusion speedup of approximately 25 % was found for a membrane consisting of palmitoyloleoylphosphatidylcholine (POPC) lipids. Crucially, the integration was stable with the 2 fs time step usually applied in all-atom MD simulations.^4^

In principle, the concept behind the atomistic TIP3P-F model – accelerating sampling by reducing the mass of the solvent – should also be applicable to coarse-grained (CG) force fields that employ explicit solvation models. Structural coarse-graining reduces the number of degrees of freedom by mapping groups of atoms onto interaction sites (“beads”). In addition to the reduced computational cost due to the smaller number of particles, CG simulations enable larger integration time steps and, in general, have faster correlation times due to their smoother (less rugged) energy landscapes. This results in a substantial sampling acceleration compared to atomistic simulations. Among the biomolecular CG force fields, Martini stands out as a widely used, general-purpose framework with a broad (bio)chemical coverage for proteins, lipids, nucleic acids, and small molecules.^12–14^ The 2021 release of Martini 3 introduced an expanded bead typology, recalibrated interaction balance, and improved transferability across molecular families, thereby improving the description of many biomolecular systems.^13^ In the Martini force field, the water model is a single Lennard-Jones particle with a mass of 72 amu, that is, four atomistic water molecules (each with a mass of 18 amu) are described by one CG bead.

Here, we develop and evaluate a fast water model that accelerates sampling in MD simulations with the Martini CG force field. To enhance the dynamics, we first systematically reduce the water bead mass and assess integration stability and simulation accuracy in NVE simulations (Section II A). To illustrate its use for biomolecular systems, we then apply the resulting *light Martini water* model to simulations of a diverse set of intrinsically disordered proteins (IDPs), spanning a range of sequence lengths and properties (Table I), and to a lipid bilayer (Section II B). We show that *light Martini water* substantially accelerates sampling without affecting equilibrium properties.

**TABLE I.** Single-chain IDP properties and simulation setups. “Length” is the number of residues of the polypeptide, “NCPR” indicates the net charge per residue and “FCR” the fraction of charged residues. “Salt” is the NaCl concentration in the simulation box. “Prod.” specifies the production simulation time.

| IDP | Length (residues) | NCPR | FCR | Salt (mM) | Box volume (nm<sup>3</sup>) | Prod. (μs) |
| --- | --- | --- | --- | --- | --- | --- |
| Hst5 | 24 | +0.21 | 0.38 | 150 | 1818 | 10 |
| Aβ40 <sup>#</sup> | 40 | -0.08 | 0.23 | 150 | 4850 (38 800) | 10 |
| A1LCD | 137<sup>‡</sup> | +0.06 | 0.12 | 50 | 9177 | 10 |
| Asyn <sup>†</sup> | 140 | -0.06 | 0.28 | 200 | 9898 | 15 |
| FUSLCD <sup>†</sup>, <sup>#</sup> | 163 | -0.01 | 0.01 | 100 | 9177 (45 255) | 10 |

<sup>†</sup>These IDPs were additionally simulated in the absence of salt (only using the minimal number of Na<sup>+</sup> ions needed to neutralize the simulation box). <sup>‡</sup>Includes an N-terminal GS tag. <sup>#</sup> These IDPs were additionally simulated in
larger simulation boxes, whose volumes are provided in parentheses.

## II. RESULTS AND DISCUSSION

### A. Development and validation of *light Martini water*

To evaluate how far the mass of the Martini water bead can be reduced without sacrificing the 20 fs integration time step used in the Martini force field, we systematically scaled down the mass and analyzed energy drifts and fluctuations in NVE simulations. The variation of the drift of the total energy (Figure 1A) indicates only minor effects down to *m_W_* = 20 amu, which is the last data point before a pronounced increase observed at lower masses. For masses < 20 amu, the energy drifts increase strongly. A similar behavior was observed for the fluctuations of the total, kinetic and potential energies (Figure 1B). For water masses ≥ 20 amu, the fluctuations of the total energy were close to zero, whereas the fluctuations of kinetic and potential energy varied around approximately 41 J/mol/particle, thereby fulfilling the expectation that the anticorrelated fluctuations of *E_pot_* and *E_kin_* are much larger than the fluctuations in *E_tot_*and cancel each other. For water masses below 20 amu, all fluctuations increase. However, the total energy fluctuations increase more strongly than those of the kinetic and potential energies, indicating increasing inaccuracies in the numerical integration of the equations of motion. While the guideline that the fluctuations of the total energy should be at least five times smaller than the fluctuations of the potential or kinetic energies^15^ are still met even for masses smaller than 20 amu, we proceeded with a mass of 20 amu for the *light Martini water* model. However, as is also shown below, numerical integration was stable also with smaller water masses.

**FIG. 1.**
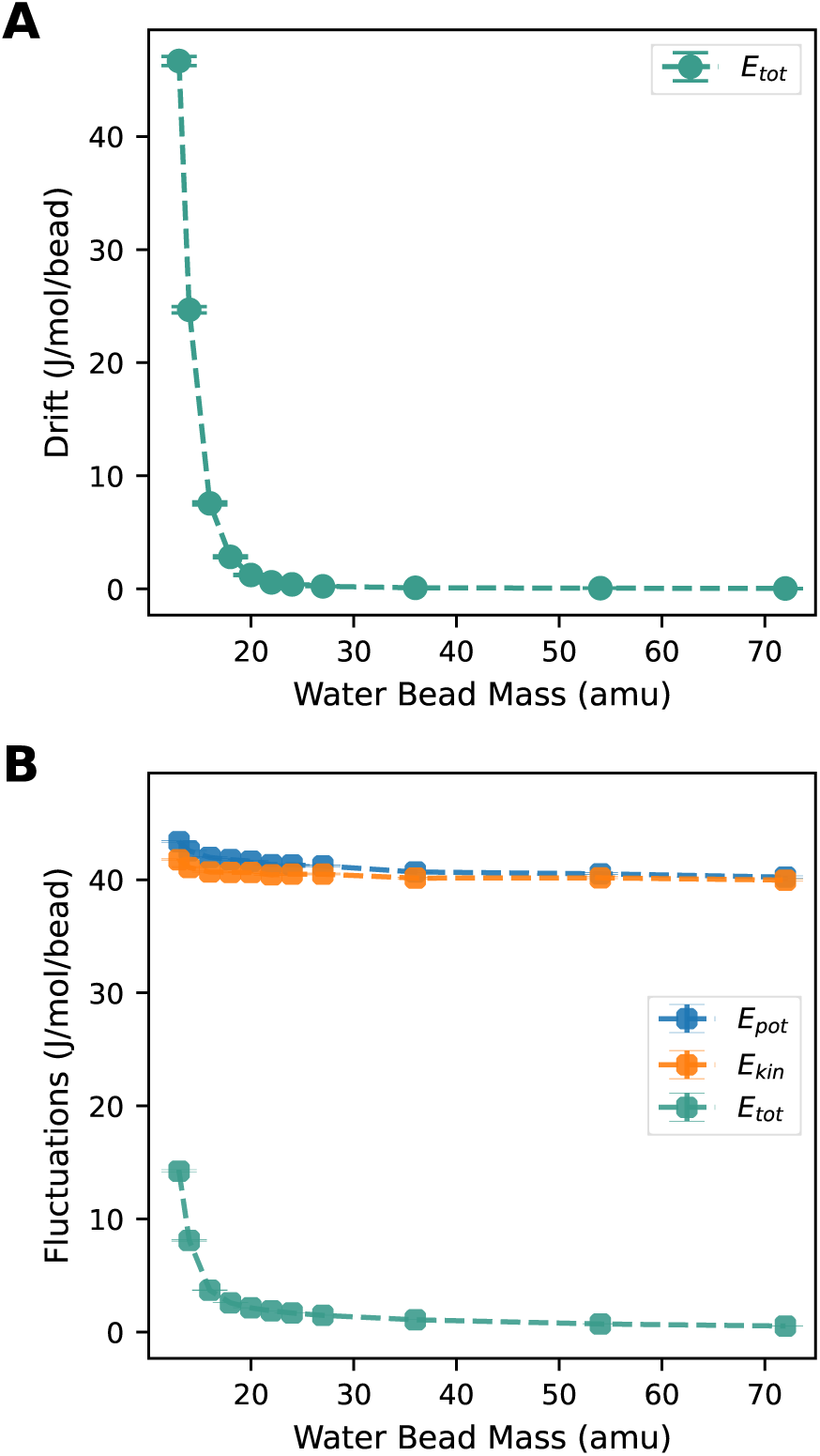
Drift of total energy over 10 ps simulation time (**A**) and fluctuations of total, kinetic and potential energies (**B**) during NVE simulations. Quantities are averages of 1000 simulation trajectories of 10 ps each. Error bars represent standard error of the mean (SEM). ployed here: The Martini water mass was reduced by a factor of 3.6 (72 to 20 amu), compared to a 16-fold mass reduction for TIP3P-F (from 18.0154 to 1.116 amu).^4^

Next, we assessed by how much decreasing the water mass accelerates dynamics in bulk water simulations. To that end, we calculated water translational self-diffusion coefficients and viscosities of bulk Martini water. The mass dependence of these quantities matches the expectations: the lower the water mass, the lower the viscosity and the higher the diffusion coefficient (Figure 2). The larger displacements of the particles per time step upon decreasing the water mass from 72 to 20 amu led a 1.8-fold faster diffusion, from (2.23 ± 0.03) to (3.96 ± 0.04) nm^2^ ns^−1^, and a correspondingly lower viscosity (from 0.66 to 0.36 mPas). The observed factor of 1.8 is close to the expected value of 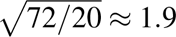, based on the relation that mass scaling by a factor *f* corresponds to time scaling by a factor √*f*. Compared to the atomistic TIP3P-F model, the observed self-diffusion speedup is of similar magnitude but more modest (1.8 versus 3.35 for TIP3P-F^4^). This comparison should, however, be interpreted in the context of the smaller mass reduction em-Having found that reducing the mass of the Martini water bead to 20 amu accelerates water dynamics without significantly compromising energy conservation, we next investigated whether the equilibrium properties of Martini water are preserved. To that end, we calculated the water bead number density as a function of bead mass and compared the radial distribution functions (RDFs) of normal (*m_W_* = 72 amu) and *light Martini water* (*m_W_* = 20 amu). The densities (Table II) span a very narrow range from 8.273 nm^−3^ (for 72 amu, corresponding to a mass density of 989.1 kgm^−3^) over 8.281 nm^−3^ (20 amu) to 8.287 nm^−3^ (14 amu). Thus, the fractional difference is 0.17 % between the largest and the smallest masses used, and 0.10 % between 72 and 20 amu. Although the data show a systematic increase in density with decreasing mass, we consider this minor difference to be negligible for practical applications.

**FIG. 2.**
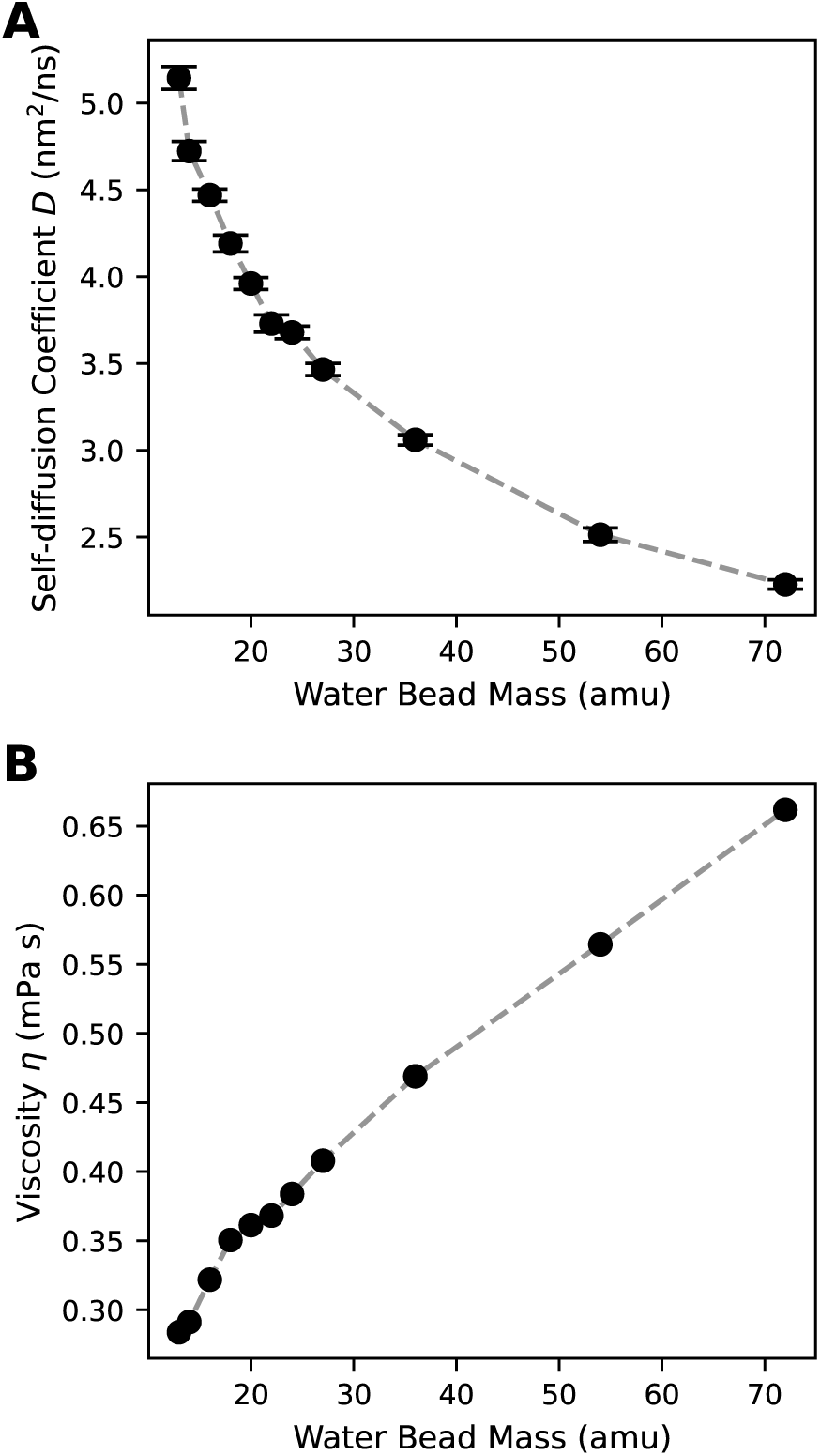
Water self-diffusion (**A**) and viscosity (**B**) as functions of water bead mass. Self-diffusion coefficients were calculated from 10 NVE simulations per system, each 100 ps long. Error bars denote standard error of the mean (SEM). Finite box-size effects were corrected as suggested by Yeh and Hummer^16^, applying the actual viscosity of the corresponding system (**B**). Viscosities were calculated from the pressure fluctuations in 50 ns NVT trajectories.

**TABLE II.** Water bead number densities from NpT simulations at 298.15 K and 1 bar. Values are means ± standard error of the mean (SEM).

| $m_W$ (amu) | $n$ (nm$^{-3}$) |
| --- | --- |
| 13 | $8.286 \pm 0.009$ |
| 14 | $8.287 \pm 0.009$ |
| 16 | $8.285 \pm 0.009$ |
| 18 | $8.285 \pm 0.009$ |
| 20 | $8.281 \pm 0.010$ |
| 22 | $8.281 \pm 0.009$ |
| 24 | $8.279 \pm 0.010$ |
| 27 | $8.279 \pm 0.009$ |
| 36 | $8.277 \pm 0.009$ |
| 54 | $8.275 \pm 0.009$ |
| 72 | $8.273 \pm 0.009$ |

The RDFs calculated for normal (72 amu) and *light Martini water* (20 amu) visually overlap on the plotted scale (Figure 3). The differences between the two curves reveal subtle deviations, which are somewhat more pronounced at the positions of the first and second peaks, with the first peak higher in the RDF of *light water*, and the second higher in the RDF of normal water (Figure 3, inset). However, the absolute differences are below 0.02, which we consider to be negligible. Therefore, we conclude that structural properties of bulk Martini water are preserved upon reduction of the bead mass to 20 amu. Taken together, our results support the choice of *m*_W_ = 20 amu as a robust setting for *light Martini water*.

**FIG. 3.**
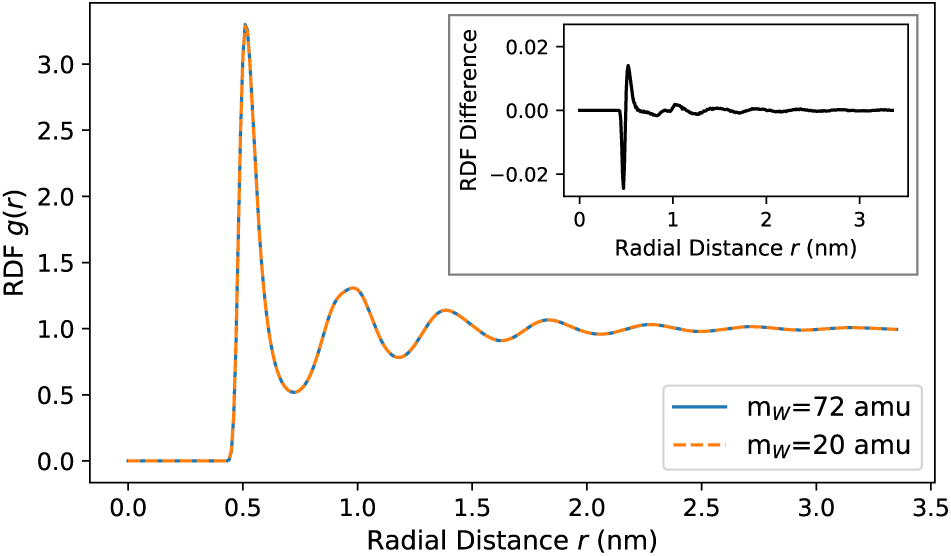
Water–water radial distribution functions (RDFs) of normal (*m_W_*= 72 amu) and *light Martini water* (*m_W_* = 20 amu). The RDFs were averaged across 1000 NVE simulation trajectories with a length of 10 ps each. The difference between the RDFs (72 amu − 20 amu) is shown in the inset.

### B. *Light Martini water* accelerates sampling of IDP conformations and lipid diffusion

#### 1. IDPs

Based on the finding that *light Martini water* has lower viscosity and faster self-diffusion, we expected that its use would accelerate the dynamics of biomolecules, at least for motions that are governed by solvent interactions. To test this hypothesis, we investigated the chain reconfiguration dynamics of a set of IDPs (Table I) and evaluated the decay of time correlation functions (TCFs) of the radius of gyration *R_g_* and end-to-end distance *R_ee_*.

The TCFs are shown for Amyloid-*β* 40 (A*β* 40) as an example in Figure 4; the data for the other IDPs are presented in the supplementary information (Figures S1, S2). For analysis, each trajectory was divided into five blocks, and a TCF was calculated for each block. The TCFs were then averaged, and a biexponential function (Equation 4, Section IV C) was fitted to the average TCF. For all systems, the fitted functions describe the data very well, with excellent agreement in the well-sampled regions of steep decay and slightly worse agreement in the noisier tails (Figures 4, S1, S2). As a quantitative measure for sampling efficiency, we used the ratio of the correlation times, *τ*_72_/*τ*_20_, where *τ* is the lag time at which the fitted function reaches 1/*e*. The resulting speedup factors are listed in Table III. Depending on the IDP and the evaluated quantity (*R_g_* or *R_ee_*), the speedup factors range from 1.08 (FUSLCD, *R_g_* TCF decay) to 2.68 (A1LCD, *R_ee_* TCF decay). In addition, to further explore how the use of *light water* accelerates sampling, we calculated diffusion coefficients of the IDPs in water (Figure 5). The speedup, defined as *D*_20_/*D*_72_, ranges from 1.48 (Asyn and FUSLCD) to 1.82 (A*β* 40 and Asyn, nosalt). Possible reasons and implications of differences between the observed speedup factors are discussed further below.

**FIG. 4.**
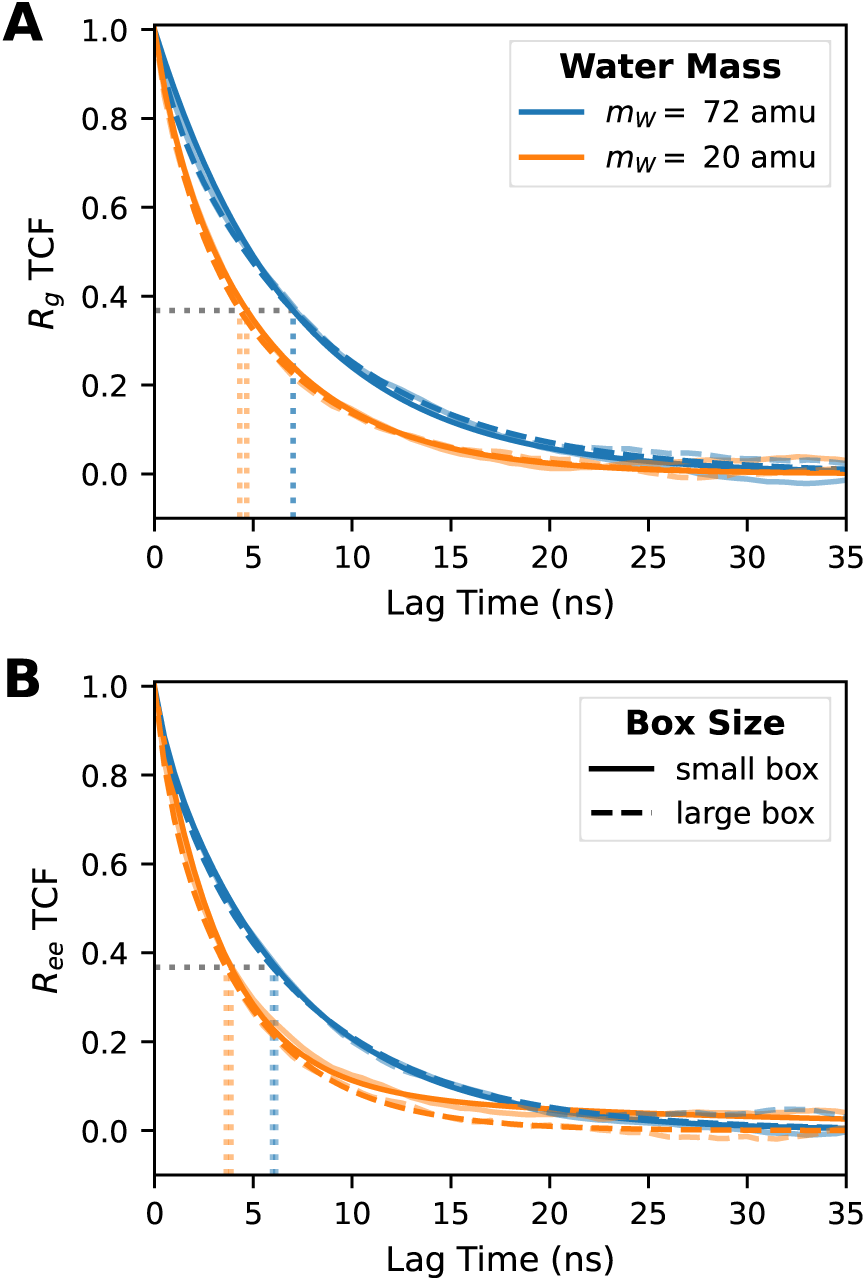
Time correlation functions of radius of gyration (**A**) and end-to-end distance (**B**) of the A*β* 40 conformations in normal (blue curves) and *light Martini water* (orange curves). The transparent curves show the raw TCFs (obtained via block averaging), whereas the opaque curves show biexponential functions fitted to the average TCFs. The vertical dashed lines indicate the lag time at which the fit function reaches 1/*e*. Two sets of simulations were performed using different box sizes with volumes of 4850 nm^3^ (“small box”, solid lines) and 38800 nm^3^ (“large box”, dashed lines), respectively. The corresponding plots for the other investigated IDPs are given in Figures S1, S2 (supplementary information).

**FIG. 5.**
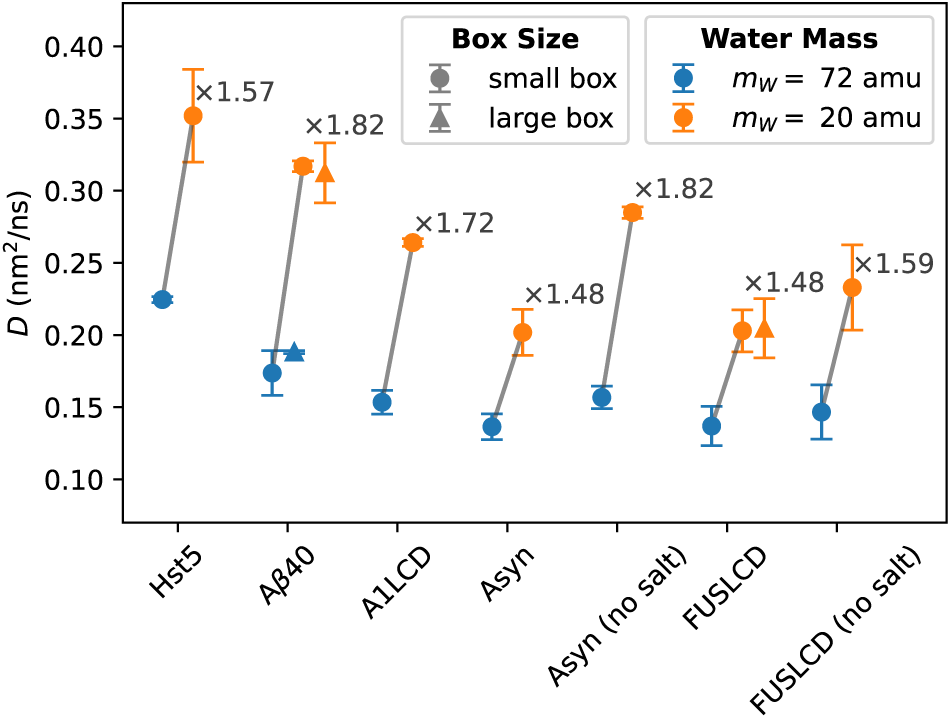
Protein diffusion coefficients *D* for normal (*m_W_*= 72 amu, blue data points) and *light Martini water* (*m_W_* = 20 amu, orange data points). The speedups (*D*_20_/*D*_72_) are given near the 20 amu data points. Finite box-size effects were corrected as suggested by Yeh and Hummer^16^, applying the viscosity of the corresponding system (see Figure 2B). For A*β* 40 and FUSLCD, diffusion coefficients obtained with larger simulation boxes are additionally depicted.

**TABLE III.** Speedup factors for radius of gyration (*R_g_*) and end-to-end distance (*R_ee_*) in *light Martini water* (*m_W_* = 20 amu) compared to normal Martini water (*m_W_* = 72 amu). The speedup factor is the ratio of the correlation times, *τ*_72_/ *τ*_20_. The underlying *τ* values are given in Table S-I (supplementary information).

| System | speedup factor |  |
| --- | --- | --- |
| | $R_g$ | $R_{ee}$ |
| Hst5 | 1.65 | 1.75 |
| Aβ40 | 1.50 | 1.59 |
| Aβ40 (large box) | 1.63 | 1.64 |
| A1LCD | 2.14 | 2.68 |
| Asyn | 1.60 | 1.57 |
| Asyn (no salt) | 1.49 | 1.69 |
| FUSLCD | 1.08 | 1.59 |
| FUSLCD (no salt) | 1.95 | 2.59 |

The sampling speedup is only meaningful if structural and thermodynamic equilibrium properties remain unaffected by the use of *light water*. This is demonstrated above for water (Figure 3), but to verify this also for the proteins, we analyzed mean values and distributions of *R_g_* and *R_ee_*. Average values and uncertainties are given in Table IV, distributions are shown in Figures 6 (for A*β* 40) and S3, S4 (supplementary information, other IDPs). The average values of *R_g_* and *R_ee_* obtained with normal and *light water* are either identical, consistent within their respective uncertainties, or only slightly shifted. The small shifts observed can plausibly be explained by finite sampling. Importantly, no mass-dependent trend was observed – *light Martini water* did not lead to consistently larger or smaller *R_g_*or *R_ee_* – supporting the conclusion that lowering the solvent mass accelerates dynamics without affecting the equilibrium distributions.

**FIG. 6.**
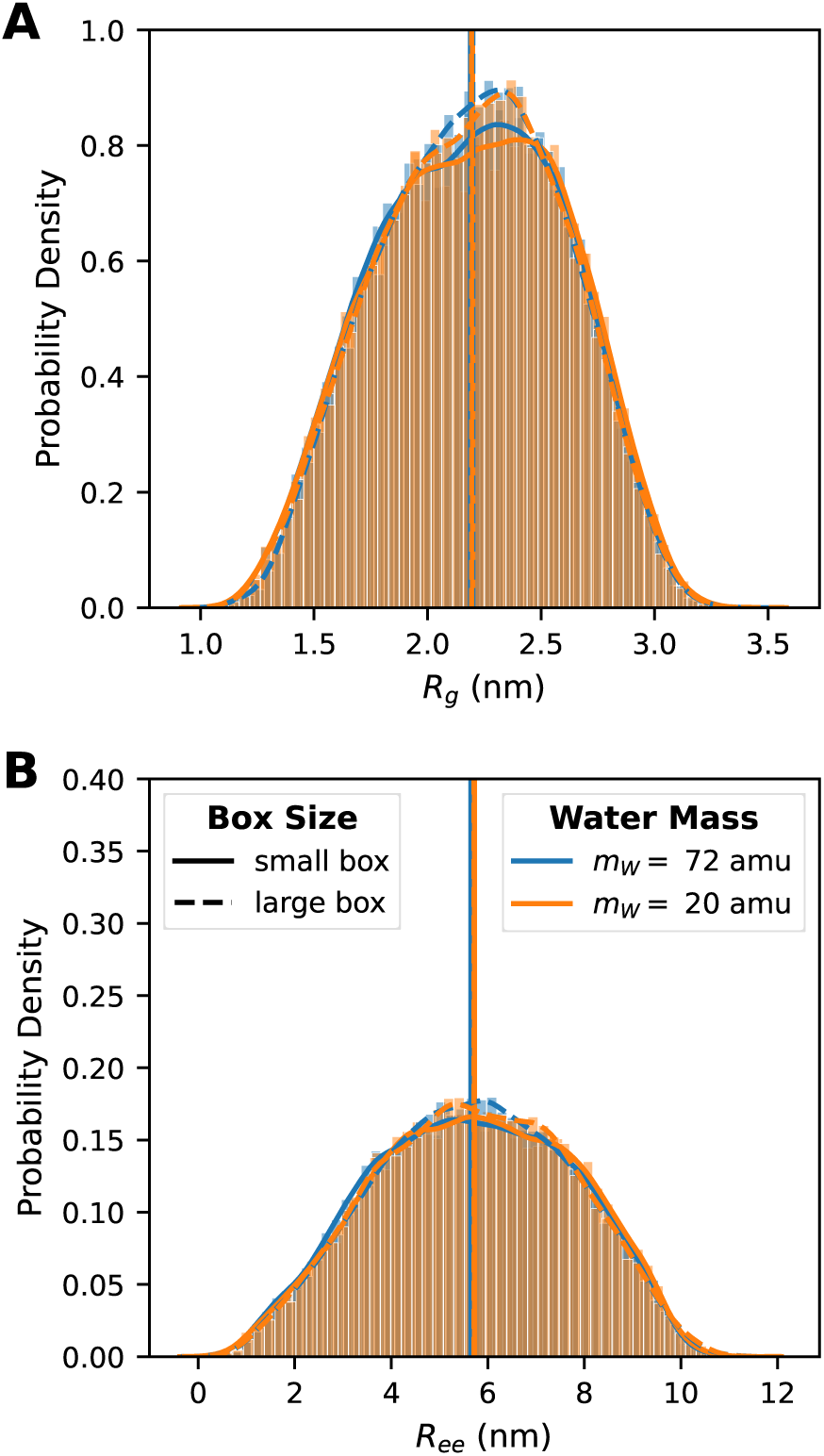
Normalized probability distributions of A*β* 40 radius of gyration (*R_g_*, **A**) and end-to-end distance (*R_ee_*, **B**) in normal (*m_W_*= 72 amu) and *light Martini water* (*m_W_* = 20 amu). The vertical lines indicate average values. The analyses were performed using differently sized simulation boxes with volumes of 4850 nm^3^ (“small box”, solid lines) and 38800 nm^3^ (“large box”, dashed lines), respectively. Equivalent plots for the other investigated IDPs are given in Figures S3, S4 (supplementary information).

**TABLE IV.** Radius of gyration (*R_g_*) and end-to-end distance (*R_ee_*) averages of single-chain IDP conformations. Given are the averages and SEM obtained from blocking. For reference, the column “comp. ref.” reports the values obtained by the Martini3-IDP developers.^17^

| System | $R_g$ (nm) | | | $R_{ee}$ (nm) | |
| --- | --- | --- | --- | --- | --- |
| | $m_W = 72$ amu | $m_W = 20$ amu | comp. ref.<sup>17</sup> | $m_W = 72$ amu | $m_W = 20$ amu |
| Hst5 | $1.62 \pm 0.01$ | $1.62 \pm 0.00$ | $1.62 \pm 0.01$ | $4.46 \pm 0.04$ | $4.45 \pm 0.03$ |
| Aβ40 | $2.19 \pm 0.02$ | $2.20 \pm 0.01$ | – | $5.65 \pm 0.08$ | $5.72 \pm 0.06$ |
| Aβ40 (large box) | $2.20 \pm 0.02$ | $2.19 \pm 0.01$ | – | $5.71 \pm 0.08$ | $5.71 \pm 0.06$ |
| A1LCD | $2.43 \pm 0.08$ | $2.30 \pm 0.04$ | $2.35 \pm 0.06$ | $4.97 \pm 0.28$ | $4.67 \pm 0.19$ |
| Asyn | $2.78 \pm 0.05$ | $2.77 \pm 0.04$ | – | $6.49 \pm 0.23$ | $6.45 \pm 0.15$ |
| Asyn (no salt) | $2.07 \pm 0.12$ | $2.03 \pm 0.03$ | – | $3.78 \pm 0.63$ | $3.60 \pm 0.12$ |
| FUSLCD | $2.83 \pm 0.05$ | $2.69 \pm 0.05$ | $2.80 \pm 0.07$ | $6.79 \pm 0.18$ | $6.22 \pm 0.17$ |
| FUSLCD (large box) | – | $2.60 \pm 0.12$ | – | – | $6.10 \pm 0.53$ |
| FUSLCD (no salt) | $5.11 \pm 0.15$ | $5.07 \pm 0.17$ | – | $10.49 \pm 0.31$ | $10.86 \pm 0.27$ |

Following the above overview in terms of sampling speedup and simulation accuracy, we next turn to a more detailed discussion of differences between the investigated IDPs. To elucidate the influence of electrostatic interactions on the conformational ensembles and the associated chain reconfiguration dynamics, two IDPs containing different fractions of charged residues (FUSLCD and Asyn, see Table I) were additionally simulated without salt (i.e., using only the minimum number of Na^+^ ions needed to neutralize the simulation box). For FUSLCD, removing salt increased ⟨*R_g_*⟩ and ⟨*R_ee_*⟩ compared to the values at 100 mM NaCl (Table IV), consistent with weaker screening and thus stronger electrostatic repulsion between the two aspartates D5 and D46, which are the only charged residues in the FUSLCD sequence. For Asyn, the simulations in the absence of salt yielded smaller ⟨*R_g_*⟩ and ⟨*R_ee_*⟩ than at 200 mM NaCl. This opposite response can be rationalized by the differences between the Asyn and FUSLCD sequences with respect to charged residues (Table I). In contrast to FUSLCD, Asyn harbors 24 negatively and 15 positively charged residues and thus has a higher fraction of charged residues. At low ionic strength, reduced screening enhances not only repulsive interactions between like charges but also attractive interactions between oppositely charged regions, such as the positively charged N-terminal and the negatively charged C-terminal parts of Asyn, thereby promoting chain compaction. At higher ionic strength, these effectively attractive interactions are more strongly screened, resulting in chain expansion.^18,19^

Regarding *R_g_*, sampling was accelerated for Hst5, Aß40, A1LCD and Asyn (200 mM salt) with speedup factors between 1.50 (A*β* 40) and 2.14 (A1LCD). In contrast, no substantial *R_g_* sampling acceleration was observed for FUSLCD at 100 mM salt (speedup factor of 1.08), suggesting that in this case, chain reconfiguration dynamics are more strongly governed by intraprotein interactions than by solvent-related friction. Interestingly, for the end-to-end distance *R_ee_*, a more pronounced speedup of 1.59 is observed (Table III), suggesting that the balance between the intramolecular and solvent-related interactions affects the decorrelation of *R_g_*and *R_ee_* in slightly different ways, with *R_ee_* being slightly more susceptible to the solvent for all IDPs studied except Asyn (Table III). While salt removal increased the sampling speedup for FUSLCD (*R_g_* speedup factor 1.08 at 100 mM salt versus 1.95 without salt), the opposite applies to Asyn (*R_g_* speedup factor 1.60 at 200 mM salt versus 1.49 without salt). These trends align with the observed salt effects on ⟨*R_g_*⟩ and ⟨*R_ee_*⟩, as discussed above: removing salt led to more compact conformations of Asyn, hence increasing the importance of intramolecular interactions relative to interactions with the solvent. In contrast, the FUSLCD ensemble shifted towards more extended conformations in the absence of salt, which are more strongly solvated. Thus, the dynamic interconversions underlying *R_g_* and *R_ee_*decorrelation are more susceptible to the lower solvent viscosity.

As controls, to examine the influence of the box size, we additionally simulated Aß40 (both in normal and *light Martini water*) and FUSLCD (only in *light Martini water*) in substantially larger simulation boxes. Results for *R_g_* and *R_ee_* sampling speedup factors, diffusion coefficients and, *R_g_* and *R_ee_* averages and distributions are presented in Tables III, IV and Figures 4, 5, 6. There were no significant differences between the quantities sampled in the simulations with the small and large simulation boxes. Additionally, all trajectories were checked for contacts with periodic images (Figures S6 S8). We conclude that the simulation boxes were sufficiently large.

In summary, our findings suggest that for Hst5, Aß40, A1LCD and Asyn, chain reconfiguration dynamics are more strongly dominated by solvent interactions compared to FUSLCD, where intrachain interactions appear to play a more important role.

#### 2. Lipid bilayers

In addition to IDPs, we applied *light Martini water* in simulations of a POPC lipid bilayer to investigate by how much the low-viscosity water enhances lipid diffusion. The lateral diffusion coefficients are (44.9±0.6) and (51.9±0.6) ·10^−12^ m^2^ s^−1^ for normal and *light water*, respectively, corresponding to a speedup factor of 1.16 (Table V). This acceleration is similar to the factor of 1.25 achieved with the TIP3P-F water model in atomistic simulations of a POPC bilayer.^4^ A plausible explanation for the smaller speedup, compared to the diffusion speedup of IDP chains in water, is that the diffusivity of lipids in a membrane is governed more by friction due to lipid–lipid interactions in the bilayer and less by friction between lipids and water. Nevertheless, the diffusion speedup obtained is still significant. In line with this interpretation, no substantial speedup was observed for a more ordered DPPC bilayer containing 30 % cholesterol (Table V), which has increased lipid packing and is thus more strongly dominated by lipid–lipid than by lipid–solvent interactions.

**TABLE V.** Lateral lipid diffusion coefficient *D_L_*, area per lipid *APL* and area compressibility modulus *K_A_* of Martini lipid bilayers (POPC and DPPC/cholesterol) in normal (*m*_W_ = 72 amu) and *light Martini water* (*m*_W_ = 20 amu). The quantities were obtained as averages ± SEM over ten NpT simulation trajectories per system, each 500 ns long. The speedup factor was calculated as *D_L_*(*m_W_* = 20 amu)/ *D_L_*(*m_W_* = 72 amu).

|  | POPC |  |  | DPPC/cholesterol |  |  |
| --- | --- | --- | --- | --- | --- | --- |
| | $m_W = 72 \text{ amu}$ | $m_W = 20 \text{ amu}$ | factor | $m_W = 72 \text{ amu}$ | $m_W = 20 \text{ amu}$ | factor |
| $D_L (\times 10^{-12} \text{ m}^2 \text{ s}^{-1})$ | $44.9 \pm 0.6$ | $51.9 \pm 0.6$ | 1.16 | $38.7 \pm 0.7$ | $39.1 \pm 0.6$ | 1.01 |
| $APL (\text{\AA}^2)$ | $58.77 \pm 0.01$ | $58.71 \pm 0.02$ | - | $43.36 \pm 0.01$ | $43.34 \pm 0.01$ | - |
| $K_A (\text{mN m}^{-1})$ | $195 \pm 4$ | $207 \pm 7$ | - | $833 \pm 7$ | $831 \pm 8$ | - |

Analogous to our workflow for the IDPs, we next examined whether structural properties of the membrane are compromised by the use of *light Martini water*. To that end, we calculated the average area per lipid (*APL*) and area compressibility modulus (*K_A_*), average density profiles (Figure 7) and order parameters of the bonds in the coarse-grained POPC lipids (Figure 8). Although there are minor differences between the respective *APL* and *K_A_* mean values (Table V), we consider them negligible. The density profiles agree well, with no apparent differences at the plotted scale (Figure 7). The absolute differences between the order parameters are below 0.002 (Figure 8, inset). Most order parameters are slightly higher for the simulations employing *light water*, suggesting marginally increased ordering; however, these differences are negligibly small. Taken together, using *light Martini water* in simulations of a POPC bilayer accelerates lipid diffusion by a factor of 1.16 without notably compromising structural properties of the membrane.

**FIG. 7.**
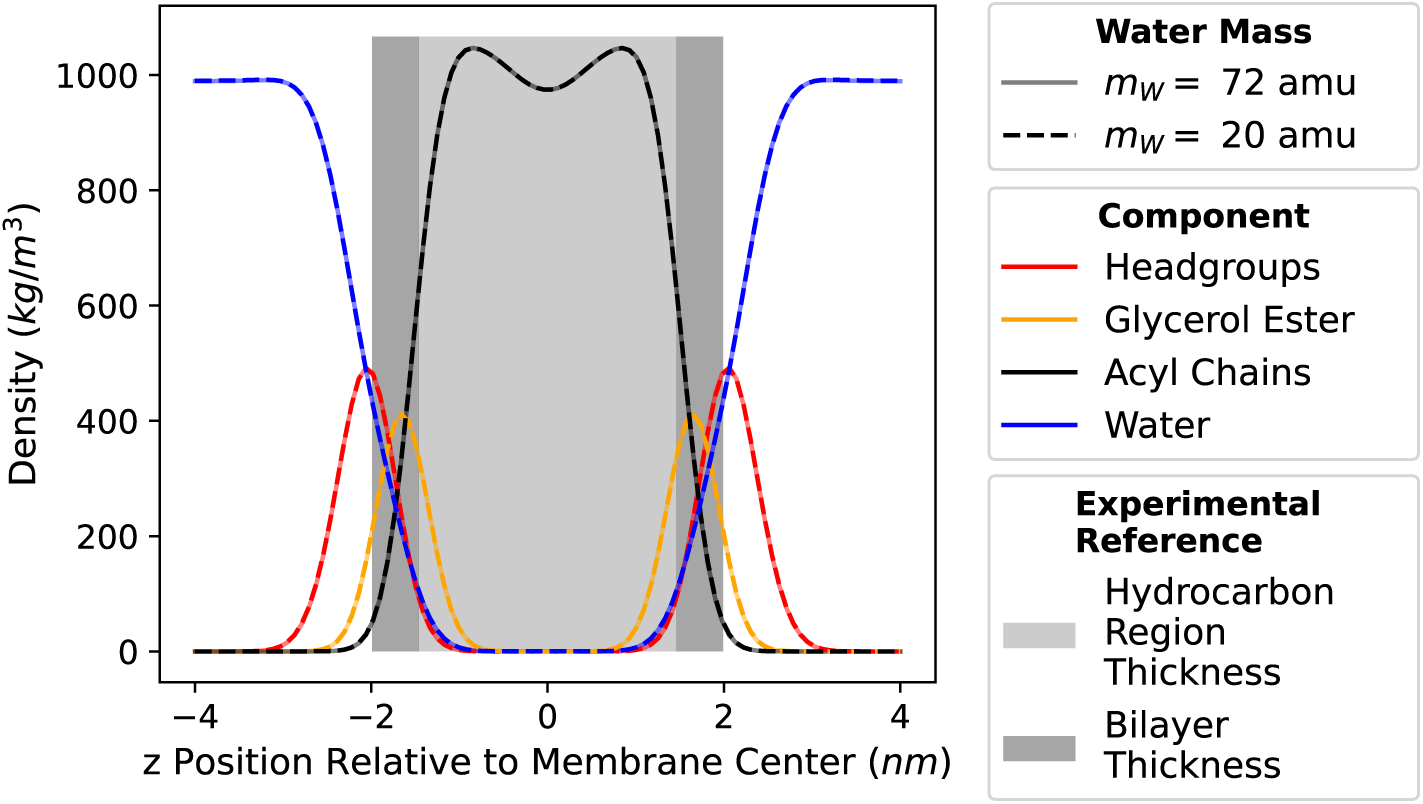
Average density profiles of a POPC bilayer in normal (*m*_W_ = 72 amu) and *light Martini water* (*m*_W_ = 20 amu). The partial mass density of the indicated components is plotted relative to the bilayer midplane. The water density of *light Martini water* was rescaled by a factor of 72/20. Experimental ranges^20^ for the hydrocarbon region thickness and bilayer thickness are illustrated in grey.

**FIG. 8.**
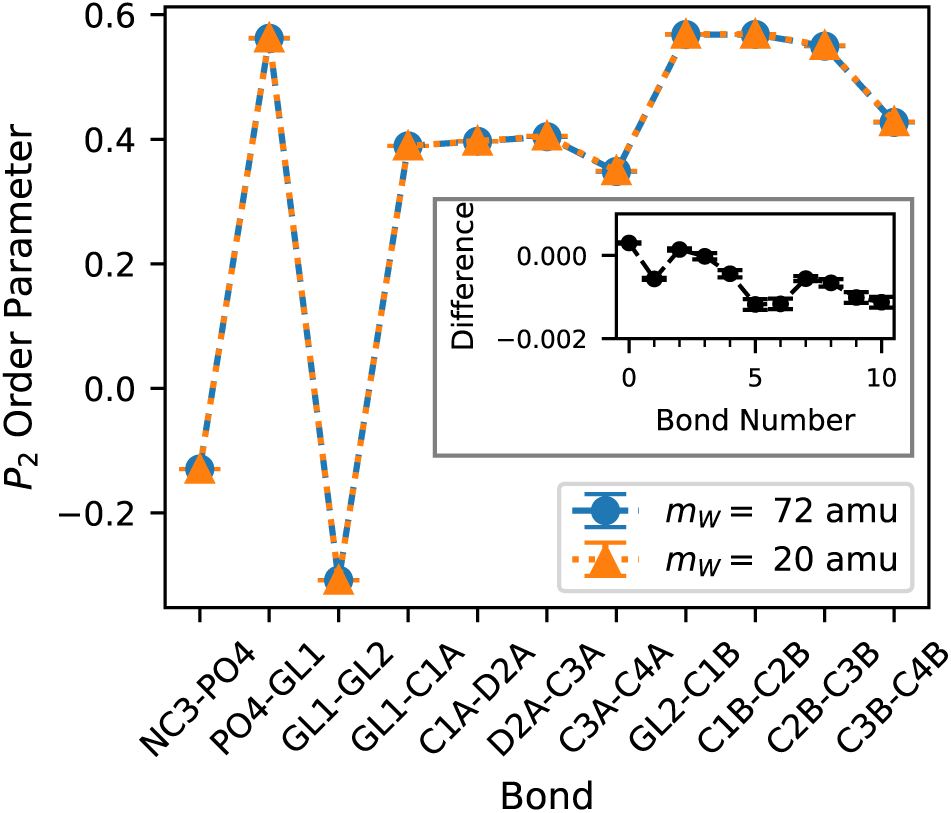
Lipid P_2_ order parameters of a POPC lipid bilayer in normal (*m*_W_ = 72 amu) and *light Martini water* (*m*_W_ = 20 amu). The values are averages over ten NpT simulation repeats of 500 ns each; error bars denote standard error of the mean (SEM). The inset shows differences between the order parameters obtained in normal and *light Martini water* (*P*_2_(*m*_W_ = 72 amu) − *P*_2_(*m*_W_ = 20 amu)).

## III. CONCLUSIONS

To accelerate sampling in coarse-grained Martini simulations using a low-viscosity water model, we systematically reduced the Martini water bead mass and assessed integration stability by monitoring energy conservation and fluctuations of the total, kinetic, and potential energies in NVE simulations. These analyses identified 20 amu as a conservative and robust choice that preserves stable integration of the equations of motion with the 20 fs time steps typically used in Martini simulations. In bulk water, this *light Martini water* has reduced viscosity and increased self-diffusion by a factor of 1.8. To determine whether these faster solvent dynamics translate into enhanced solute dynamics, we applied the model to two distinct classes of biomolecular systems: IDPs and lipid bilayers. As quantified by the decorrelation of *R_g_* and *R_ee_*, *light Martini water* accelerated IDP chain reconfiguration by up to a factor of 2.68, depending on the protein, while translational diffusion increased by up to a factor of 1.82. Lateral lipid diffusion exhibited a smaller, yet still significant, speedup of 1.16 for a POPC lipid bilayer. Importantly, these gains in sampling efficiency were achieved without affecting equilibrium properties, as demonstrated by water densities and RDFs, IDP *R_g_* and *R_ee_* distributions, and the area per lipid, area compressibility modulus, density profile, and lipid order parameters.

Based on these findings, we propose *light Martini water* as a versatile water model that accelerates sampling in coarse-grained Martini simulations. The model is trivial to implement, does not introduce any computational overhead, and is expected to be broadly applicable in all simulations employing the Martini 3 force field and its derivatives. The exact sampling speedup achieved with *light Martini water* depends on the particular system and dynamic mode under investigation, and the sampling acceleration might even be zero (speedup factor 1.0), as demonstrated in this work for the DPPC/cholesterol bilayer. However, *light Martini water* should never slow down dynamics.

Of course, the “native” dynamics of the original Martini model will be perturbed in *light Martini water*. However, for most applications this is not a major concern, because time scales from coarsegrained simulations generally require system- and observable-specific mapping for comparison with physical time scales (e.g., from experiments or all-atom simulations).^21,22^ In general, beyond accelerating the conformational sampling of biomolecules, *light Martini water* may be particularly useful for simulating processes in which slow diffusive dynamics are important and represent a sampling bottleneck, such as IDP condensate formation/phase separation or the sampling of protein–protein and protein–ligand interactions. Exploring these and other applications, as well as extending the concept to other solvents, are possible directions for future research.

## IV. METHODS

### A. General settings

The standard Martini 3 force field^13^ was used in bulk water simulations, the Martini 3-IDP force field^17^ in IDP simulations and the Martini 3 lipidome parameters^23^ in the lipid bilayer simulations. Unless stated otherwise, simulations were performed as follows, using GROMACS 2025.2.^24^ Energy minimizations were performed with the steepest-descent algorithm until convergence to machine precision. The equations of motion were solved with the leap-frog integrator, using a time step of 20 fs and employing periodic boundary conditions. For the neighbor search, the Verlet cutoff scheme with an 1.35 nm cutoff and an update frequency of every 20 steps was used, with disabled automated buffer setting, following the recommendations of Kim et al.^25^ For electrostatics, a relative dielectric constant of 15 was used together with a reaction-field approach with a cutoff of 1.1 nm and an infinite reaction-field dielectric (epsilon_rf = 0). Lennard-Jones (van der Waals) interactions were treated with a cutoff at 1.1 nm and the potential-shift-Verlet modifier. Bond constraints defined in the force-field topologies were solved with LINCS.^26,27^ Temperature was controlled with the v-rescale thermostat^28^ with a coupling time constant of *τ_T_* = 1 ps. For pressure control, the c-rescale barostat^29^ was used with a target pressure of 1 bar, a coupling time constant of *τ_p_* = 10 ps and a compressibility of 3 × 10^−4^ bar^−1^.

Uncertainties were calculated as standard error of the mean (SEM), if multiple simulation repeats were considered for calculation of a quantity, or by block averaging (`gmx analyze -ee`), if only a single trajectory was considered.

### B. Development of light Martini water

A cubic box of neat Martini water (*N* = 2588 beads) with initial box length *L* = 7 nm was built. Single precision was used for all runs, except that the NVE simulations (see below) were performed in double precision with per-step energy evaluation and neighbor list update. To probe solvent-mass effects, independent systems differing only in the water-bead mass *m*_W_ were prepared: 13, 14, 16, 18, 20, 22, 24, 27, 36, 54, and 72 amu (72 amu is the reference Martini water mass). For each system, the following procedure was conducted: After energy minimization, the system was equilibrated for 15 ns at a temperature of 298.15 K, followed by 50 ns of NpT equilibration. This trajectory was used for density analysis (Table II). A frame with box dimensions close to the average box dimensions was extracted from the NpT trajectory, and the box lengths were set to the precise average values. Starting from this frame, a 50 ns NVT run was conducted. From this trajectory, the coordinates and velocities of 1000 equally spaced frames were extracted, which served as starting points for 1000 NVE simulations, with a length of 10 ps each, using GROMACS 2018.3 in double-precision. In these NVE simulations, the velocity-Verlet integrator was used instead of leap-frog. The NVE trajectories were evaluated with respect to total energy drifts and fluctuations of total, kinetic and potential energy (Figure 1). Furthermore, the water-water RDFs (Figure 3) were calculated based on these NVE trajectories. The water self-diffusion coefficients (Figure 2A) were calculated from 10 extended NVE trajectories, each 100 ps long, using `gmx msd`. Viscosities were calculated from a 50 ns NVT simulation initiated from the configuration obtained from NpT equilibration as described above. The pressure tensor was recorded at every simulation step (every 20 fs) and used to calculate the autocorrelation functions of its nondiagonal elements and the viscosity via the Green–Kubo equation (Equation 2), which expresses the viscosity as the time integral of the pressure autocorrelation function^30,31^

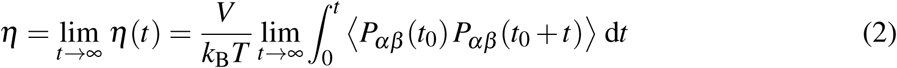

Here, *k_B_* is Boltzmann’s constant, *T* the absolute temperature, *V* the simulation box volume and *P_αβ_* are the nondiagonal pressure tensor elements. The pressure autocorrelation functions and the corresponding Green-Kubo integrals were calculated using the code^32^ developed and applied by Prass *et al*.^33^ Then, an exponential function (Equation 3)

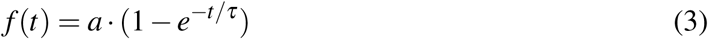

was fitted to the Green-Kubo integrals in a fit range of 0 ≤ *t* ≤ 100 ps, and the viscosity was extracted from the asymptotic limit of the fitted function.

### C. Simulation of IDPs

For each IDP chain listed in Table I, two otherwise identical systems were prepared that differed only in the water mass (normal (*m_W_* = 72 amu) and *light Martini water* (*m_W_* = 20 amu)). The initial IDP conformations were generated with AlphaFold 3^34^ and converted to the Martini3-IDP representation with martinize2.^35^ Each protein was placed in a dodecahedral box with the edge length set to the values reported for the corresponding single-chain setups in the Martini3-IDP publication.^17^ To test for box size effects, Aß40 (both water masses) and FUSLCD (only *light Martini water*) were additionally simulated in larger boxes. Boxes were filled with Martini water using `gmx solvate` and neutralized; NaCl was then added with `gmx genion` to reach the target ion concentration. Each system was energy-minimized, followed by a 15 ns NVT equilibration (target temperature 303 K) and a 50 ns NpT equilibration (target pressure 1 bar). During these equilibrations, harmonic position restraints were applied to the backbone beads. Production MD simulations were then run in the NpT ensemble, with the total simulation times provided in Table I. The radius of gyration *R_g_* was calculated with `gmx gyrate`, taking into account all protein beads for analysis. The end-to-end distance *R_ee_*, defined as the distance between the first and last backbone beads, was obtained with `gmx distance`. Mean values and uncertainties were obtained by blocking analysis with `gmx analyze -ee`. For *R_g_*(*t*) and *R_ee_*(*t*), normalized time correlation functions were computed with `gmx analyze -ac`. To that end, each production trajectory was partitioned into 5 equal, non-overlapping blocks. A biexponential function (Equation 4)

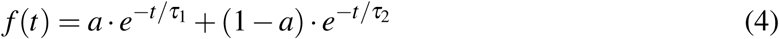

was fitted to the averaged TCF over a system-specific lag time window ([0,*t*_fit_], with *t*_fit_ set to 20, 35, 200, 200 and 200 ns for Hst5, Aß40, A1LCD, Asyn and FUSLCD, respectively). The TCF decay (correlation) time *τ* was defined as the lag time at which the fit function reaches 1/*e*. In addition, we took the TCF(*τ*) = 1/*e* values directly from the raw TCFs, as model free estimates. The relative deviation between the two decay times is below 15 %, confirming the good agreement between the fitted functions and the raw data and the robustness of the estimated *τ* values.

To analyze protein translational diffusion, the mean squared displacement (MSD) of the IDP center of mass was analyzed in the production NpT simulations using `gmx msd` and the diffusion coefficients *D* were obtained from Equation 5 by a linear fit to the MSD as a function of lag time *t*,

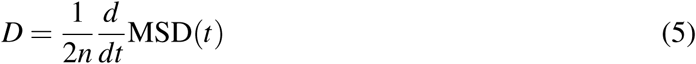

where 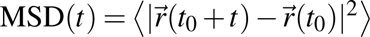 is the mean squared displacement of the particle position (or of the center of mass of a group of particles), and *n* denotes the number of spatial dimensions considered, i.e., *n* = 3 for diffusion in three-dimensional space (and *n* = 2 for the lateral diffusion in the membrane plane, see below). For the IDPs, the MSD(*t*) curves are inherently noisy for longer lag times, because the analysis is carried out for a single particle (or single center of mass). Thus, to focus the analysis on a lag time regime with good statistics, the linear fit was performed over the lag time window of 20 to 100 ns, that is, only up to 1 % of the trajectory length.^36–38^ The underlying MSD curves are shown in Figure S5 (supplementary information). Uncertainties were estimated by dividing the fitting range into two subranges (20 to 60 ns and 60 to 100 ns) and performing linear fits for each subrange. The absolute difference between the resulting diffusion coefficients was used as an error estimate. To account for finite box-size effects, the correction suggested by Yeh and Hummer^16^ was applied, using the constant *ξ* = 4.58486 for rhombic dodecahedrons proposed by Cao et al.^39^ and the viscosity of the corresponding system (see Figure 2B). The speedup in diffusion was quantified as the ratio *D*_20_/*D*_72_.

### D. Simulation of lipid bilayers

The following procedure was applied for a lipid bilayer comprised of 1024 POPC molecules (512 per leaflet) as well as for a lipid bilayer comprised of 1024 dipalmitoylphosphatidylcholine (DPPC) molecules (512 per leaflet) and 438 cholesterol molecules (219 per leaflet, amounting to 30 mol-%). The coarse-grained simulation boxes of the lipid bilayers in water were generated using the *Martini Maker* tool^40,41^ in CHARMM-GUI.^42^ As the refined Martini3 lipid parameters, recently published by Pedersen et al.,^43^ were not implemented in the CHARMM-GUI *Martini Maker* yet, the Martini3.0.0 force field was selected in the *Martini Maker* tool and the generated parameters were subsequently replaced by the refined lipid parameters. In one simulation system, the Martini water bead mass was changed to 20 amu. After energy minimization, the systems were equilibrated for 15 ns in the NVT ensemble (target temperature *T* = 298.15 K for the POPC membrane and 320 K for the DPPC/cholesterol membrane) and for 50 ns in the NpT ensemble (target pressure 1 bar, semi-isotropic pressure coupling scheme). Starting from different initial velocities taken from a Maxwell-Boltzmann distribution at 298.15 K for the POPC membrane and 320 K for the DPPC/cholesterol membrane, 10 NpT simulation runs of 500 ns each were carried out, saving coordinates to disk every 10 ps.

All properties mentioned in the following were calculated as averages over all production simulation repeats. To quantify the lateral lipid diffusion coefficients *D_L_*, the MSD of the phosphate head group beads in the xy-plane was calculated using `gmx msd` and the diffusion coefficients were obtained from Equation 5 by fitting a linear regression to the MSD as a function of lag time. The fit was performed over a lag-time range of 25 to 200 ns, corresponding to 5 to 40 % of the trajectory length. The underlying MSD curves are shown in Figure S9 (supplementary information).

Area per lipid *APL* and area compressibility modulus *K_A_* were calculated based on the *x* and *y* box dimensions, using equations 6 and 7, respectively.

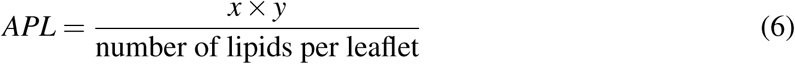

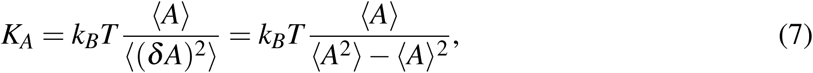

with ⟨*A*⟩ being the average area, ⟨(*δ A*)^2^⟩ = ⟨*A*^2^⟩ − ⟨*A*⟩^2^ the variance of the area, *k_B_* Boltzmann’s constant and *T* the absolute temperature.

The density profiles were calculated using `gmx density` with the -center and -symm options. For the individual components analyzed, the following Martini lipid beads were taken into account: head groups: NC3, PO4; glycerol ester: GL1, GL2; acyl chains: C1A, D2A, C3A, C4A, C1B, C2B, C3B, C4B. Coarse-grained (*P*_2_) order parameters were calculated with an in-house Python code using the following equation,

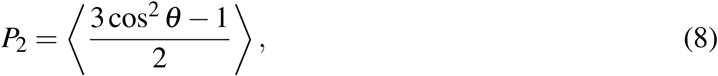

where *θ* is the angle between the bilayer normal (*z*-axis of the box) and the bond between the respective coarse-grained beads.

## V. SUPPLEMENTARY MATERIAL

The supplementary material contains all *R_g_* and *R_ee_* TCFs (Figures S1,S2) and the decorrelation times derived from them (Table S-I), *R_g_*and *R_ee_* distributions (Figures S3,S4), data on minimum distances to periodic images (Figures S6-S8) of all IDP simulation systems and the MSD curves underlying the calculation of diffusion coefficients (Figure S5 for IDPs and S9 for lipids).

## VI. DATA AVAILABILITY

The data that support the findings of this study are openly available in RESOLVdata/TUDOdata, at https://doi.org/10.17877/RESOLV-2026-MS4UJFFM.

## VII. AUTHOR DECLARATIONS

The authors declare no conflict of interest.

## VIII. AUTHOR CONTRIBUTIONS

A.E. and A.P.Z.: Conceptualization, methodology, investigation, data curation, formal analysis, visualization, writing – original draft, writing – review and editing. L.V.S.: Supervision, conceptualization, funding acquisition, validation, writing – review and editing.

## IX. ACKNOWLEDGEMENT

This work was funded by the Deutsche Forschungsgemeinschaft (DFG, German Research Foundation) under Germany’s Excellence Strategy – EXC-2033 – 390677874 – RESOLV.

