## Supplementary Information for "Light Martini water accelerates sampling in coarse-grained molecular dynamics simulations"

Table S-I: Time metrics for radius of gyration ($R_g$) and end-to-end distance ($R_{ee}$) decorrelation in normal ($m_{\mathrm{W}}$ = 72 amu) and *light* ($m_{\mathrm{W}}$ = 20 amu) Martini water. The decorrelation time $\tau$ is defined as the time at which the TCF fit function reaches $1/e$. The speedup factor was obtained by dividing the decorrelation times obtained with normal and *light* water ($\tau(m_W = 72$ amu$)\,/\,\tau(m_W = 20$ amu$)$).

| System | $R_g$ | | | $R_{ee}$ | | |
|---|---|---|---|---|---|---|
| | $\tau$ (ns) | | speedup | $\tau$ (ns) | | speedup |
| | $m_W = 72$ | $m_W = 20$ | | $m_W = 72$ | $m_W = 20$ | |
| Hst5 | 3.64 | 2.21 | 1.65 | 3.72 | 2.13 | 1.75 |
| A$\beta$40 | 7.01 | 4.67 | 1.50 | 6.14 | 3.87 | 1.59 |
| A$\beta$40 (large box) | 7.02 | 4.32 | 1.63 | 5.97 | 3.63 | 1.64 |
| A1LCD | 47.49 | 22.24 | 2.14 | 29.39 | 10.96 | 2.68 |
| Asyn | 42.78 | 26.81 | 1.60 | 31.42 | 20.03 | 1.57 |
| Asyn (no salt) | 30.21 | 20.23 | 1.49 | 26.62 | 15.78 | 1.69 |
| FUSLCD | 31.78 | 29.40 | 1.08 | 22.23 | 13.98 | 1.59 |
| FUSLCD (large box) | – | 30.57 | – | – | 16.98 | – |
| FUSLCD (no salt) | 65.25 | 33.46 | 1.95 | 30.19 | 11.65 | 2.59 |

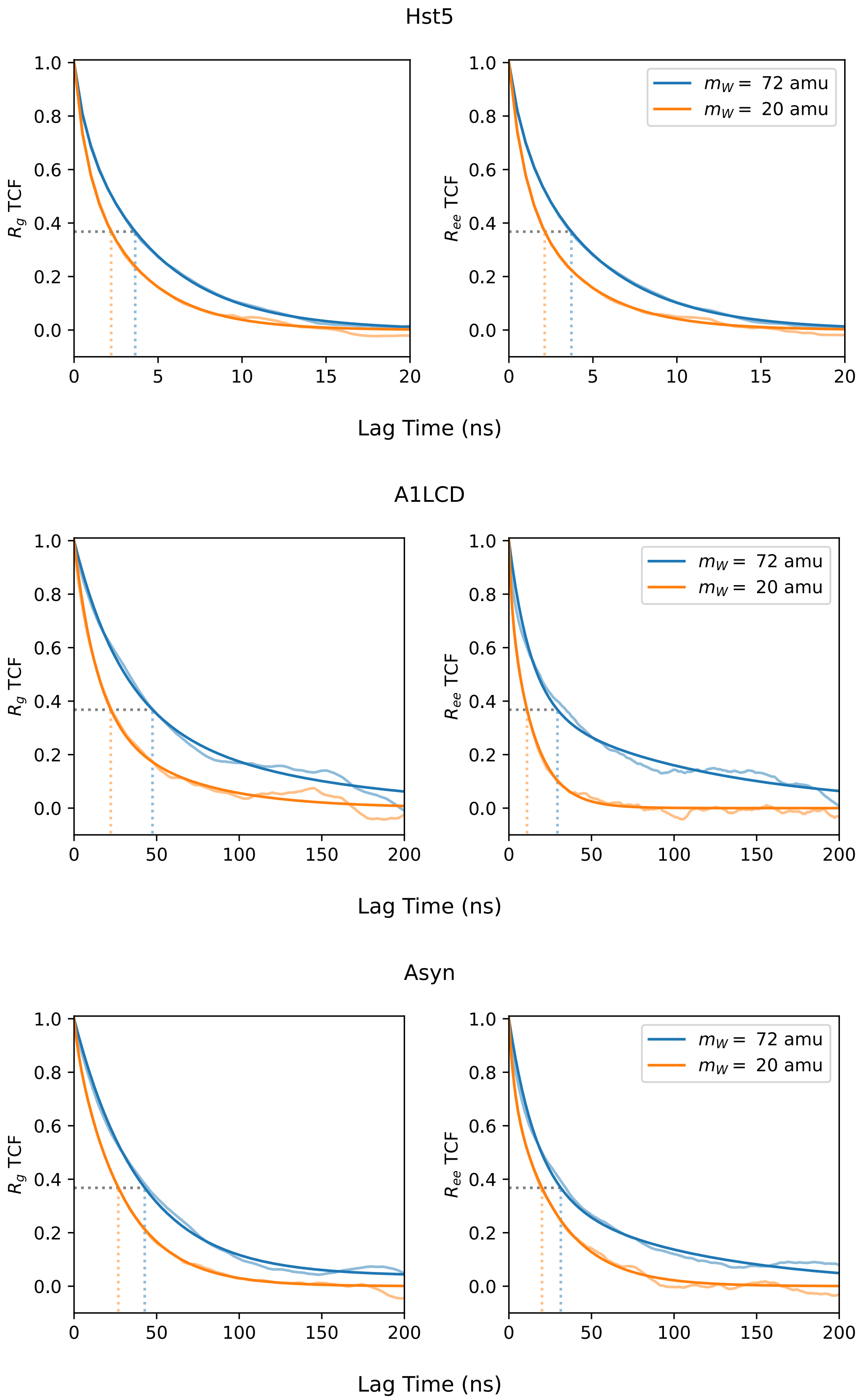


Figure S1: Time correlation functions of radius of gyration ($R_g$) and end-to-end distance ($R_{ee}$) of the Hst5, A1LCD and Asyn conformations in normal ($m_W = 72$ amu) and *light* ($m_W = 20$ amu) Martini water. The transparent curves show the averaged TCF, whereas the opaque curves show bi-exponential functions fitted to the average TCF. In each case, the whole lag time range shown was considered for the fit.

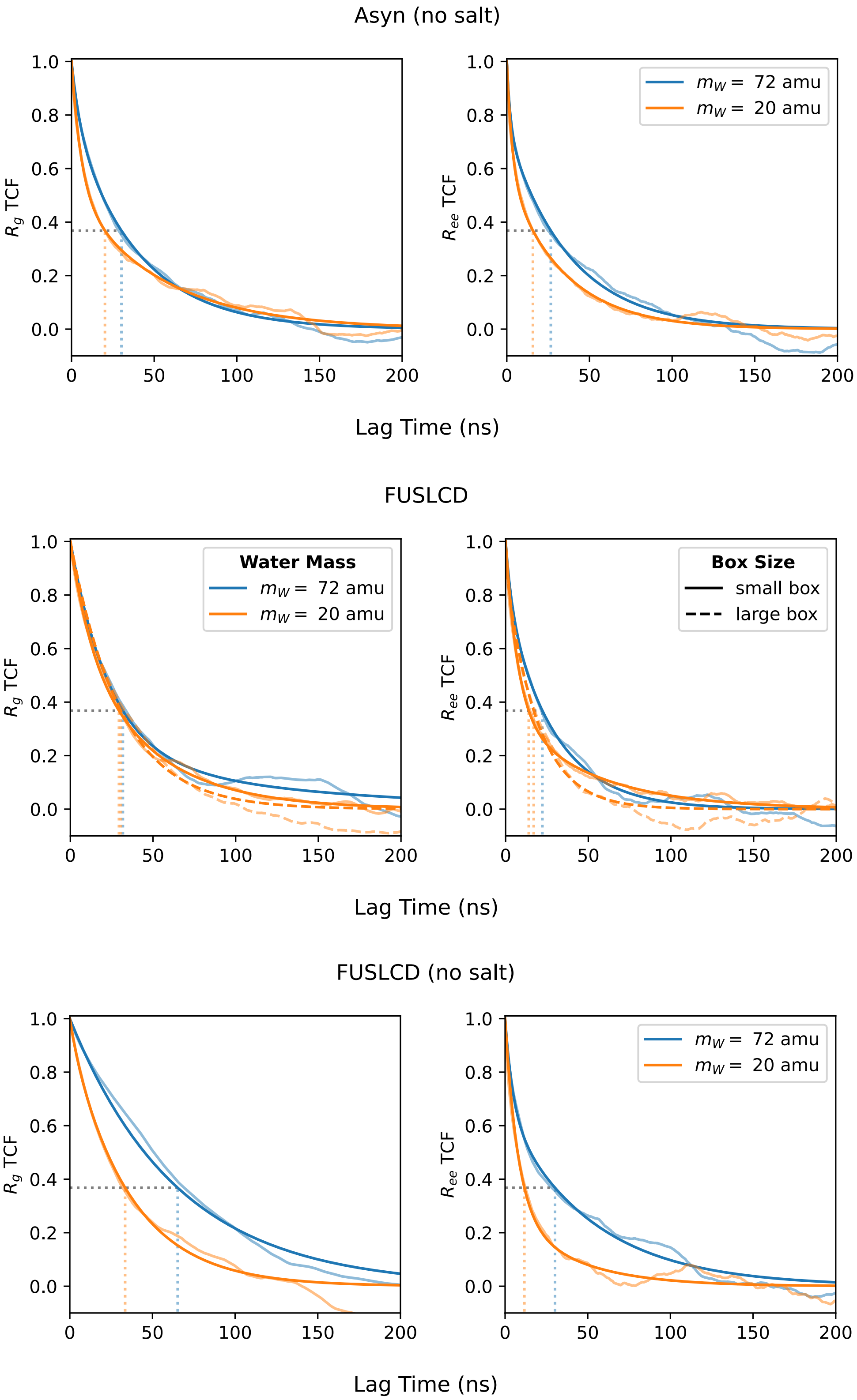


Figure S2: Time correlation functions of radius of gyration ($R_g$) and end-to-end distance ($R_{ee}$) of the Asyn (no salt), FUSLCD and FUSLCD (no salt) conformations in normal ($m_W = 72$ amu) and *light* ($m_W = 20$ amu) Martini water. The transparent curves show the averaged TCF, whereas the opaque curves show bi-exponential functions fitted to the average TCF. In each case, the whole lag time range shown was considered for the fit. For FUSLCD, these analyses were performed using differently sized dodecahedral boxes with volumes of 9177 $nm^3$ ("small box", solid lines) and 45 255 $nm^3$ ("large box", dashed lines), respectively.

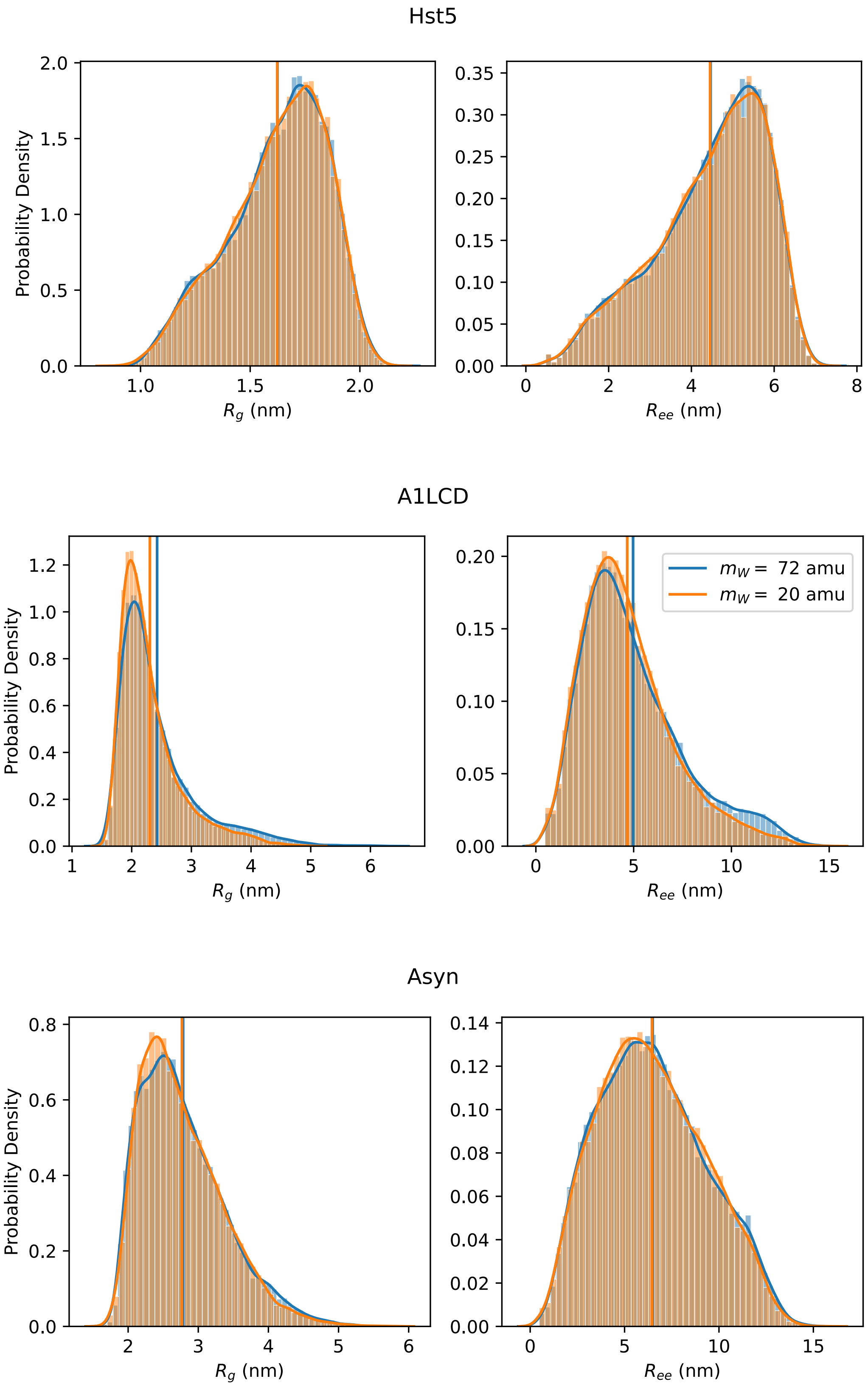


Figure S3: Normalized probability distributions of Hst5, A1LCD and Asyn radius of gyration ($R_g$, **A**) and end-to-end distance ($R_{ee}$, **B**) in normal ($m_W = 72$ amu) and *light* ($m_W = 20$ amu) Martini water. The vertical lines indicate average values.

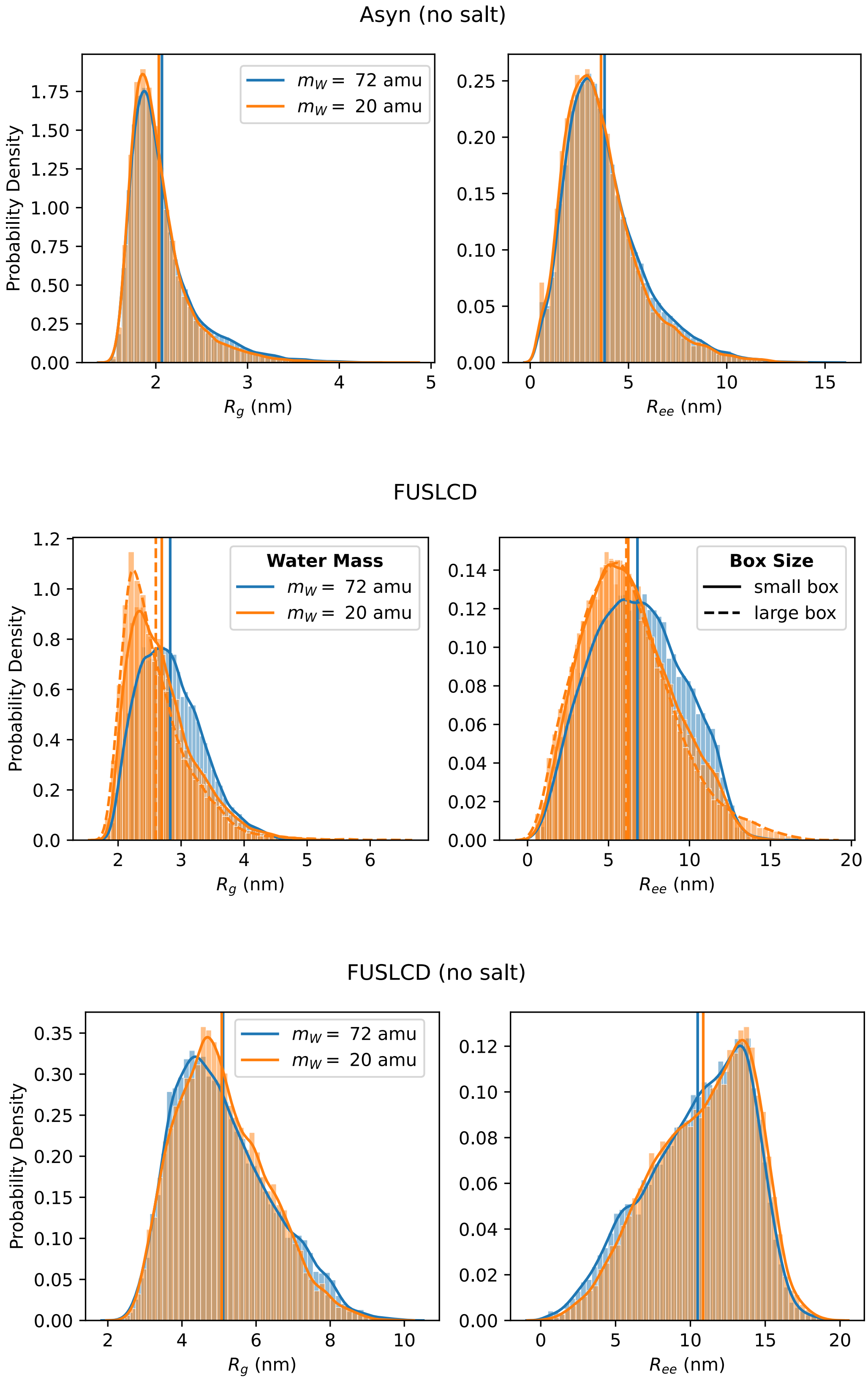


Figure S4: Normalized probability distributions of Asyn (no salt), FUSLCD and FUSLCD (no salt) radius of gyration ($R_g$, **A**) and end-to-end distance ($R_{ee}$, **B**) in normal ($m_W$ = 72 amu) and *light Martini water* ($m_W$ = 20 amu). The vertical lines indicate average values. For FUSLCD, these analyses were performed using differently sized dodecahedral boxes with volumes of 9177 $nm^3$ ("small box", solid lines) and 45 255 $nm^3$ ("large box", dashed lines), respectively.

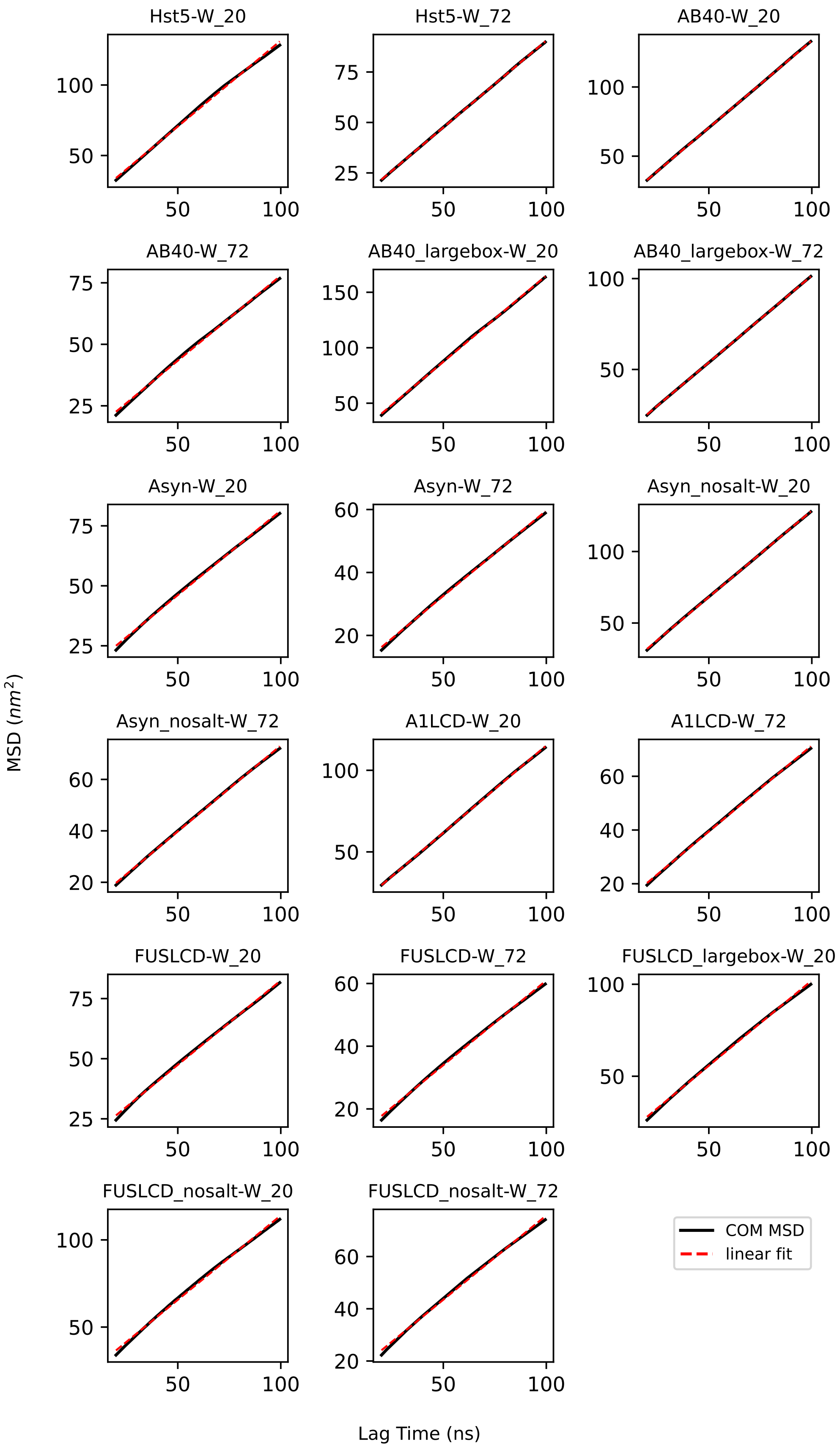


Figure S5: Center-of-mass (COM) mean-squared-displacement (MSD) plots used to calculate the translational diffusion coefficients of the investigated single-chain IDPs in standard ("W_72") and *light* ("W_20") Martini water. The red dashed lines represent the linear fit performed to calculate the diffusion coefficients.

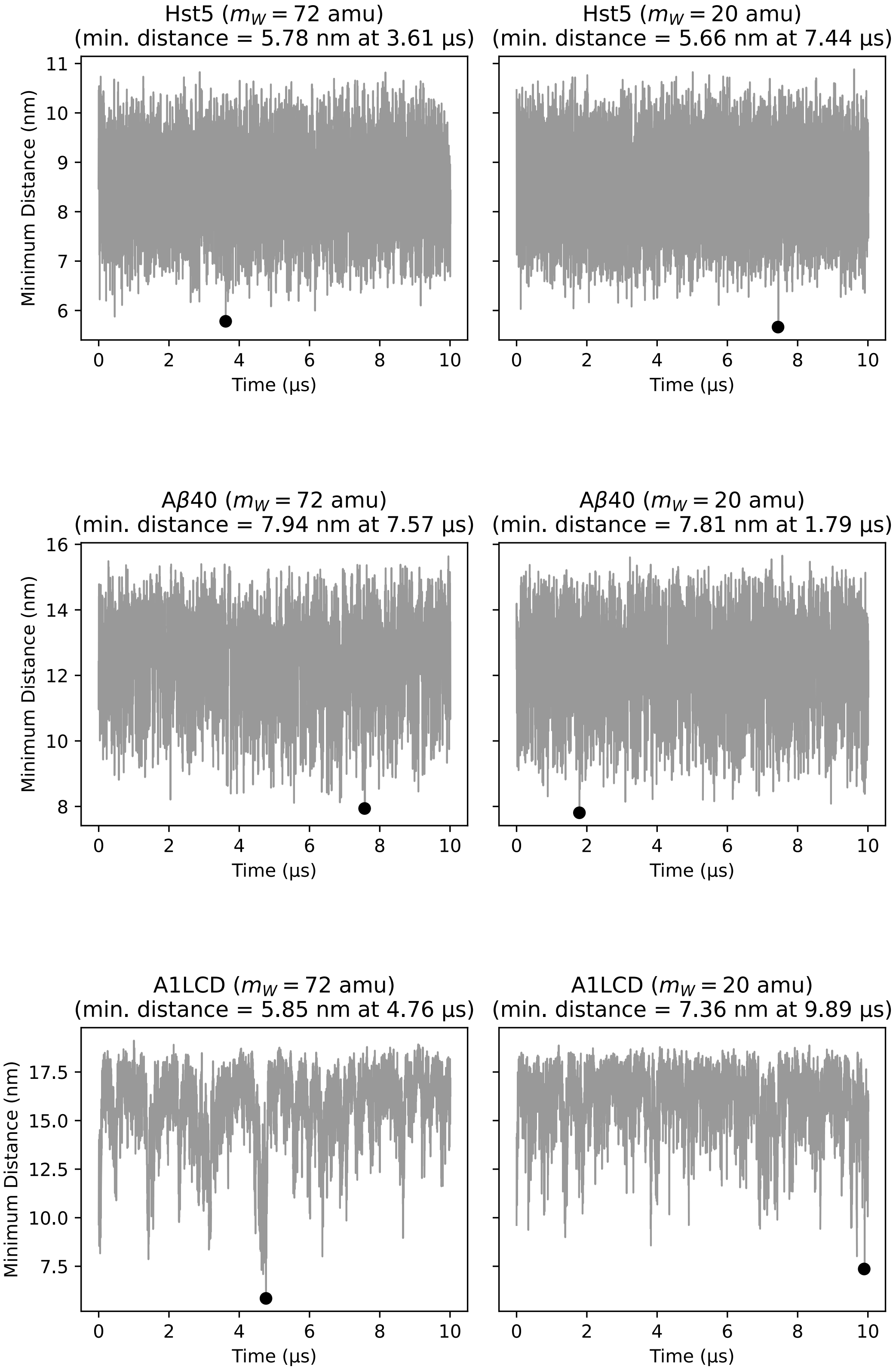


Figure S6: Minimum periodic–image distance vs. time for Hst5, Aß40, A1LCD. Each panel shows the minimal distance to the periodic image between any two protein beads in the course of the simulations. The filled marker indicates the absolute minimum; panel titles include the extracted minimum and its time from `gmx mindist`.

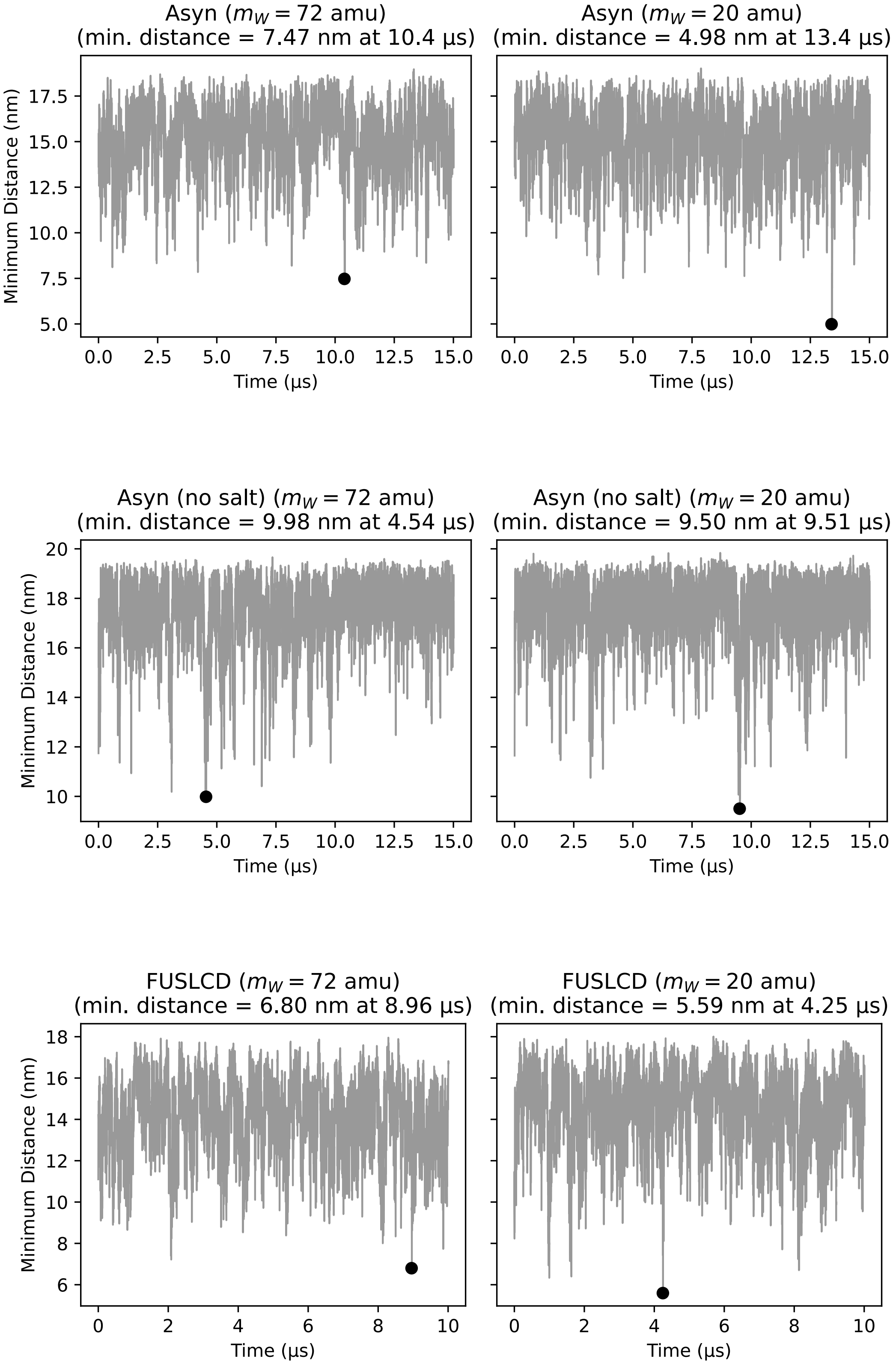


Figure S7: Minimum periodic–image distance vs. time for Asyn, Asyn (no salt), FUSLCD. Each panel shows the minimal distance to the periodic image between any two protein beads in the course of the simulations. The filled marker indicates the absolute minimum; panel titles include the extracted minimum and its time from `gmx mindist`.

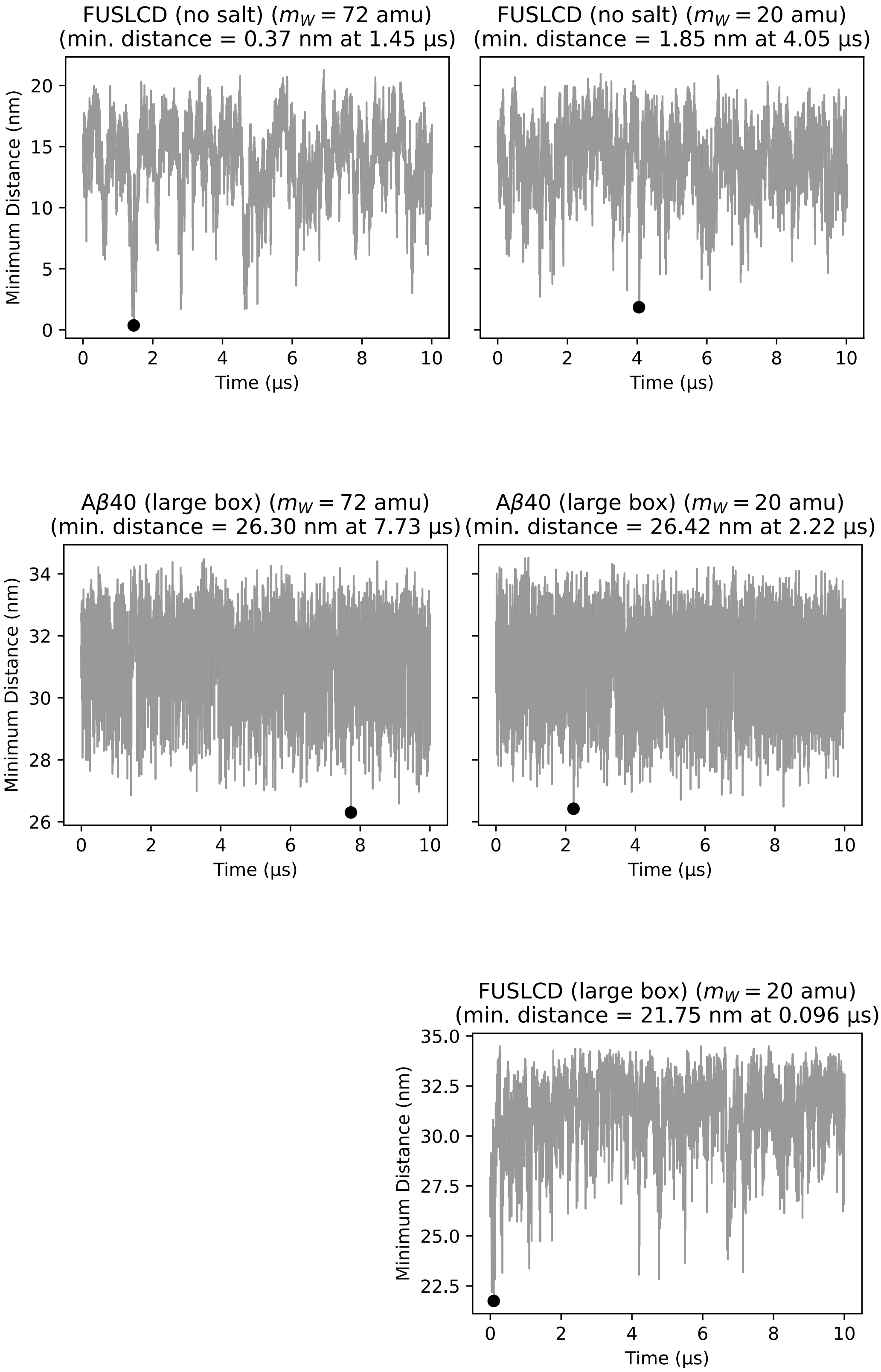


Figure S8: Minimum periodic–image distance vs. time for FUSLCD (without salt), Aß40 (large box), FUSLCD (large box, only *light Martini water*). Each panel shows the minimal distance to the periodic image between any two protein beads in the course of the simulations. The filled marker indicates the absolute minimum; panel titles include the extracted minimum and its time from `gmx mindist`.

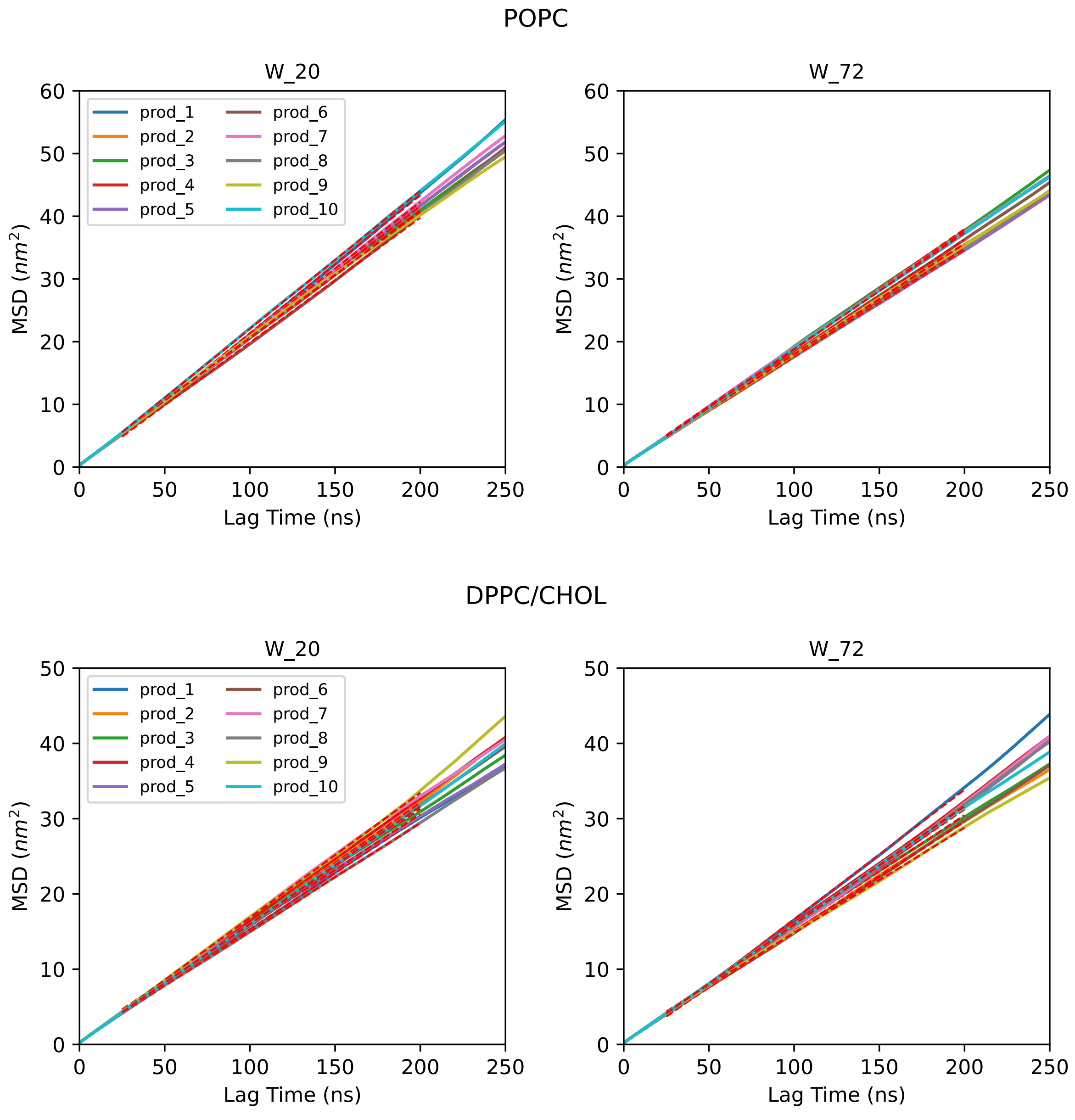


Figure S9: Mean-squared-displacement (MSD) plots used to calculate the lateral lipid diffusion coefficients of the investigated lipid bilayers in standard ("W_72") and *light* ("W_20") Martini water. Each curve represents the lateral MSD of the phosphate group beads in a single simulation trajectory. The red dashed lines represent the linear fits performed to calculate the diffusion coefficients.
